# Modulation of inhibitory communication coordinates looking and reaching

**DOI:** 10.1101/2020.06.16.156125

**Authors:** Maureen A. Hagan, Bijan Pesaran

## Abstract

Looking and reaching are controlled by different brain regions and coordinated during natural behaviour^1–3^. Understanding how flexible, natural behaviours like coordinated looking-and-reaching are controlled depends on understanding how neurons in different regions of the brain communicate^4^. Excitatory multiregional communication recruits neural coherence in a gamma-frequency (40-90 Hz) band^5^. Inhibitory control mechanisms are also required to flexibly control behaviour^6^, but little is known about how neurons in one region transiently suppress individual neurons in another to support behaviour. How does neuronal firing in a sender-region transiently suppress firing in a receiver-region? Here, we study inhibitory communication during a flexible, natural behaviour, termed gaze-anchoring, in which saccades are transiently inhibited by coordinated reaches. During gaze-anchoring, we find that neurons in the reach region of the posterior parietal cortex can inhibit neuronal firing in the parietal saccade region to suppress eye movements and improve reach accuracy. Importantly, suppression is transient, only present around the coordinated reach, and greatest when reach neurons fire spikes at a particular phase of beta-frequency (15-25 Hz) activity, not gamma-frequency activity. Our work provides evidence in the activity of single neurons for a novel mechanism of inhibitory communication in which beta-frequency neural coherence transiently inhibits multiregional communication to flexibly coordinate our natural behaviour.

## Main

The flexible control of behaviour depends on both excitatory and inhibitory mechanisms to route information flow between cortical regions^4^. Excitatory projection neurons can drive increases in activity in downstream regions and enhance stimulus processing by recruiting correlated^7^ and coherent^8–12^ temporal patterns of neural activity. Inhibitory control mechanisms also guide behaviour in the face of changing goals and contingencies^6^. However, whether and how increased firing of neurons in one cortical region can improve behavioural performance by suppressing firing in another cortical region remains poorly understood. How does inhibitory communication between neurons in different brain regions guide behaviour flexibly?

In primates, saccadic eye movements are naturally coordinated with arm movements to make accurate reach-and-grasp movements^13,14^. The reach and saccade brain systems of the posterior parietal cortex are interconnected by excitatory projections across short white matter tracts, also called U-fibers^15,16^. Silencing neural firing in the parietal reach region (PRR) alters reaching and not saccades made alone, while silencing firing in the parietal saccade region (the lateral intraparietal area, area LIP) alters saccades but not reaching^17,18^. Thus, communication between neurons in PRR and neurons in area LIP may support coordinated visual behaviour.

In humans, behavioural inhibition improves reach performance through gaze-anchoring^19^. Gaze is naturally ‘anchored’ to the target of an ongoing reach and new eye movements are inhibited, extending target foveation in time and improving reach accuracy. Reach-region neurons guiding the reach may inhibit response selection in saccade-region neurons responsible for the upcoming saccade. We therefore tested the activity of individual neurons in the parietal reach (PRR) and saccade (area LIP) regions for evidence of inhibitory communication during gaze-anchoring (**Fig 1A)**.

**Figure 1.**
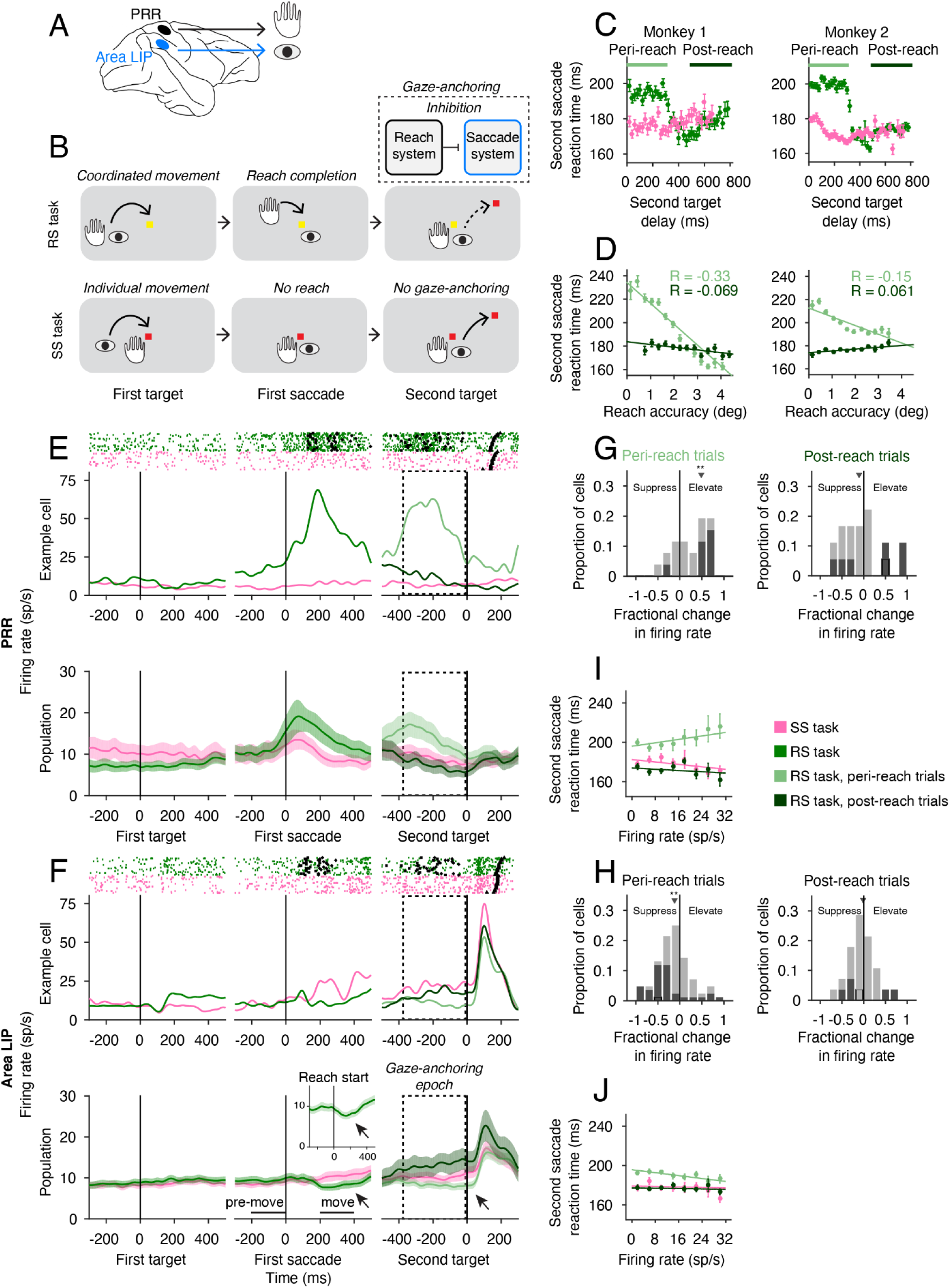
Coordinated behaviour provides evidence of inhibitory multiregional communication. **(A)** Schematic showing parietal reach region (PRR) and saccade region (lateral intraparietal area, area LIP) which guide reaching and looking, respectively and communicate during coordinated eye-hand movements. (**B**) Reach and saccade double-step task (RS) and Saccade double-step task (SS), indicating hand and eye position at each task epoch. The reach system may transiently inhibit the saccade system during gaze-anchoring to improve behavioural performance. (**C**) Reaction times for the second saccade (SSRT) as a function of second target delay after reach completion for the RS task (green) and SS task (pink). Horizontal lines indicate peri-reach (light green) and post-reach trials (dark green). (**D**) SSRT as a function of reach accuracy on peri-reach (light green) and post-reach (dark green) trials. (**E**) Example cells and population firing rates for PRR and (**F**) LIP. Inset shows LIP population data aligned to reach onset in the RS task. Arrows indicate suppression of LIP firing rates due to reach onset. Dashed box indicates the gaze anchoring epoch used as the analysis window for neural data. On rasters, reach start (squares), reach stop (diamond) and second saccade (triangle) are shown for each example trial. Dashed box indicates gaze-anchoring epoch used for all subsequent neural analysis. (**G**) Fractional change in firing rate ((RS-SS)/SS) between the two tasks for PRR and (**H**) LIP. Cells which showed significant difference (p<0.05, dark squares) and example cells (from F, G, black outline) are indicated. (**I**) Firing rates in PRR and (**J**) LIP on each trial type as a function of SSRT.

### Inhibitory communication modulates behaviour

We trained two non-human primates (*Macaca mulatta*) to perform a behavioural task that naturally elicited gaze-anchoring, the reach-saccade task, as well as a saccade-only task which should not elicit gaze-anchoring (**Fig 1B, Extended Data Fig 1**). In the reach-saccade task (RS task), each monkey made a reach and saccade to a target, followed by a second saccade to a newly-presented target. In the saccade-only task (SS task), each monkey made a sequence of two saccades and no reach. In the RS task, we presented the second saccade target after reach completion (second target delay, 0-800 ms after the reach). We matched the time interval from the first saccade to presentation of the second saccade target across the SS and RS tasks. Monkeys were not rewarded for making fast or slow eye movements in either task.

Both monkeys exhibited gaze-anchoring naturally, without training (**Methods, Fig 1C**). SSRTs were significantly longer on peri-reach trials, when the second target appeared within 300 ms of the reach, compared with post-reach trials, when the second target appeared 500-800 ms after reach completion. (**Fig 1C**, peri-reach: M1: p=5×10^−62^, 8140 trials, M2: p= 5×10^−210^, 10,245 trials; post-reach: M1: p=1×10^−3^, 2872 trials, M2: p=0.54, 3903 trials; t-test, compared to saccade trials matched for second target delay). Reach accuracy, measured as degrees of visual angle of the reach endpoint from the reach target (see **Methods**), also varied with SSRT on peri-reach but not post-reach trials (**Fig 1D**). On peri-reach trials, reaching was significantly more accurate on trials with longer SSRTs (**Fig 1D**, peri-reach: M1: R=-0.33, slope=-17.8 ms/deg, p=1×10^−95^, M2: R=-0.15, slope=-7.5 ms/deg, p=2×10^−33^, R: Pearson correlation). On post-reach trials, the association between reach accuracy and SSRT was weaker and inconsistent (**Fig 1D**, post-reach: M1: R=-0.069, slope=-2.3 ms/deg, p=2×10^−4^, M2: R=0.061, slope=1.5 ms/deg, p=6×10^−5^).

These data reveal a temporal window for gaze-anchoring around the time of reach completion. Trial-by-trial changes in the SSRT revealed a correlation with the reach reaction time in the RS task, but not with the saccade reaction time in the SS task (**Extended Data Fig 2**). Thus, gaze-anchoring occurs briefly around the reach and involves trial-by-trial changes in reach and saccade movement performance.

### Reaching inhibits saccade firing

To investigate single neuron activity for inhibitory communication during gaze-anchoring, we recorded from 120 spatially-selective neurons in the parietal reach and saccade systems (PRR - 34 neurons. Area LIP - 86 neurons; **Methods, Extended Data Fig 3**). We presented the first movement target in the response field of a PRR neuron and the second target in the response field of an area LIP neuron (**Methods**). Consistent with a role in guiding the reach, and previous work^20^, PRR neurons fired more during coordinated reaches than during saccades made alone (**Fig 1E**). Consistent with previous work suggesting a role in guiding saccades^21^, area LIP neurons fired more before the saccade when the second target was in the response field (**Fig 1F**).

We defined a gaze-anchoring epoch from 350 ms before second target onset until second target onset (dashed box). On peri-reach trials, the gaze-anchoring epoch included activity related to reach execution, reach preparation and the coordinated saccade. On post-reach trials and saccade trials, these processes were weaker or absent during the gaze-anchoring epoch.

To test for firing rate differences between the RS task and SS task, we defined a task selectivity index that measured the fractional difference (MFD) in firing rate ((RS-SS)/SS) during the gaze-anchoring epoch. Area LIP activity was transiently suppressed during RS trials around the time of the reach (Fig 1F, move vs pre-move: MFD=-0.18, p=0.01, sign-rank test) and no change was observed on SS trials (MFD=0.01, p=0.10, sign-rank test). Aligning responses to the start of the reach revealed that the suppression of area LIP activity started at the start of the reach (Fig 1F inset). Comparing RS and SS trials, PRR neurons fired significantly more around the coordinated reach and area LIP neurons fired significantly less (Fig 1G,H peri-reach: PRR: MFD=0.49, p=3.7×10^−4^. LIP: MFD=-0.11, p=0.001, sign-rank test). Firing rates did not differ 500 ms after the reach (Fig 1G,H post-reach: PRR: MFD=-0.17, p=0.24; Area LIP: MFD=-0.04, p=0.51, sign-rank test). PRR neurons fired more prior to first target onset in the SS task compared to the RS task. Since initial hand position differed between the two tasks, PRR activity before first target onset may reflect initial hand position^22,23^. Consistent with this, PRR neuron firing did not significantly differ before target onset during a center-out reach-and-saccade task and a center-out saccade-alone task when the initial eye-hand position was the same (Methods, p=0.87, permutation test). Therefore, increases in the firing of neurons in PRR may drive inhibition and suppress firing in area LIP specifically during gaze-anchoring.

Trial-by-trial changes in area LIP and PRR firing rate also reflected gaze-anchoring. On peri-reach trials, PRR neurons fired more and area LIP neurons fired less on trials with longer SSRTs compared with trials with shorter SSRTs (**Fig 1I**,**J. PRR:** R=0.11, slope=0.44 ms/(sp·s^-1^), p=0.001. **LIP:** R=-0.10, slope=-0.37 ms/(sp·s^-1^), p=7×10^−7^, Pearson correlation). This inverse relationship is transient and task-dependent: neural gaze-anchoring effects were specific to coordinated movements and were not observed at other times during the RS task (post-reach: PRR: R=-0.06, p=0.35. Area LIP: R=-0.03, p=0.2; Saccade: PRR: R=-0.12, slope=-0.31 ms/(sp·s^-1^), p=0.004. Area LIP: R=-0.02, p=0.54. Pearson correlation). Firing rates of a subset of simultaneously recorded area LIP and PRR neurons were negatively correlated during gaze anchoring trials, but positively correlated during other trials (32 pairs, peri-reach trials: R=-0.07, p=0.02; post-reach trials: R=0.2, p=5×10^−6^; saccade trials: R=0.08, p=0.01; Spearman’s correlation).

### Dual coherent beta-LFP modulates gaze anchoring

The correlations in firing rate between PRR and area LIP that we observe suggests that gaze anchoring is due to neurons in PRR communicating with neurons in area LIP. To better understand such multiregional communication, we reasoned that behavioural performance should vary with reach-to-saccade communication trial-to-trial. We therefore analyzed how performance varies with neural activity on RS task peri-reach trials compared to RS task post-reach trials and SS task saccade trials.

How might one region exert a transient, task-dependent inhibitory/suppressive effect on another? Neuronal coherence is the correlated timing of neural activity across groups of neurons measured by the phase of local field potential (LFP) activity in specific frequency bands^24^. Since the strength of neural interactions depends on the timing of neuronal activity with respect to neural excitability, multiregional communication may depend on the phase of neural coherence. Spike timing and LFP activity in the beta-frequency band reflects suppression of movement initiation^20,25,26^, motor processing^27,28^, top-down feedback^29–31^ and multiregional integration^32–34^. As a result, the phase of beta-frequency neuronal coherence may support inhibitory communication between the reach and saccade systems. If so, inhibition between PRR and area LIP, and the associated behavioural performance, should vary trial-to-trial according to the timing of spikes with respect to the phase of beta-frequency activity in PRR and area LIP.

To test the relationship between behavioural performance and the phase of beta-frequency coherence, we conducted 151 experimental sessions with PRR spiking recorded simultaneously with local field potentials (LFP) in both PRR and area LIP (**Fig 2A**). In the RS task, LFP activity in area LIP and PRR transiently synchronized around the time of the reach, with PRR spiking tending to occur when the beta-band of LFP activity in both areas was at a particular phase (Example trial: **Fig 2B**, box). Whereas on SS trials, during the same relative time period, PRR spiking tended to occur when beta-band LFP activity in both areas was at a different phase (Example trial: **Fig 2C**, box). The task-dependent change in the phase of PRR spiking was present across trials (**Extended Data Fig 4**).

**Figure 2.**
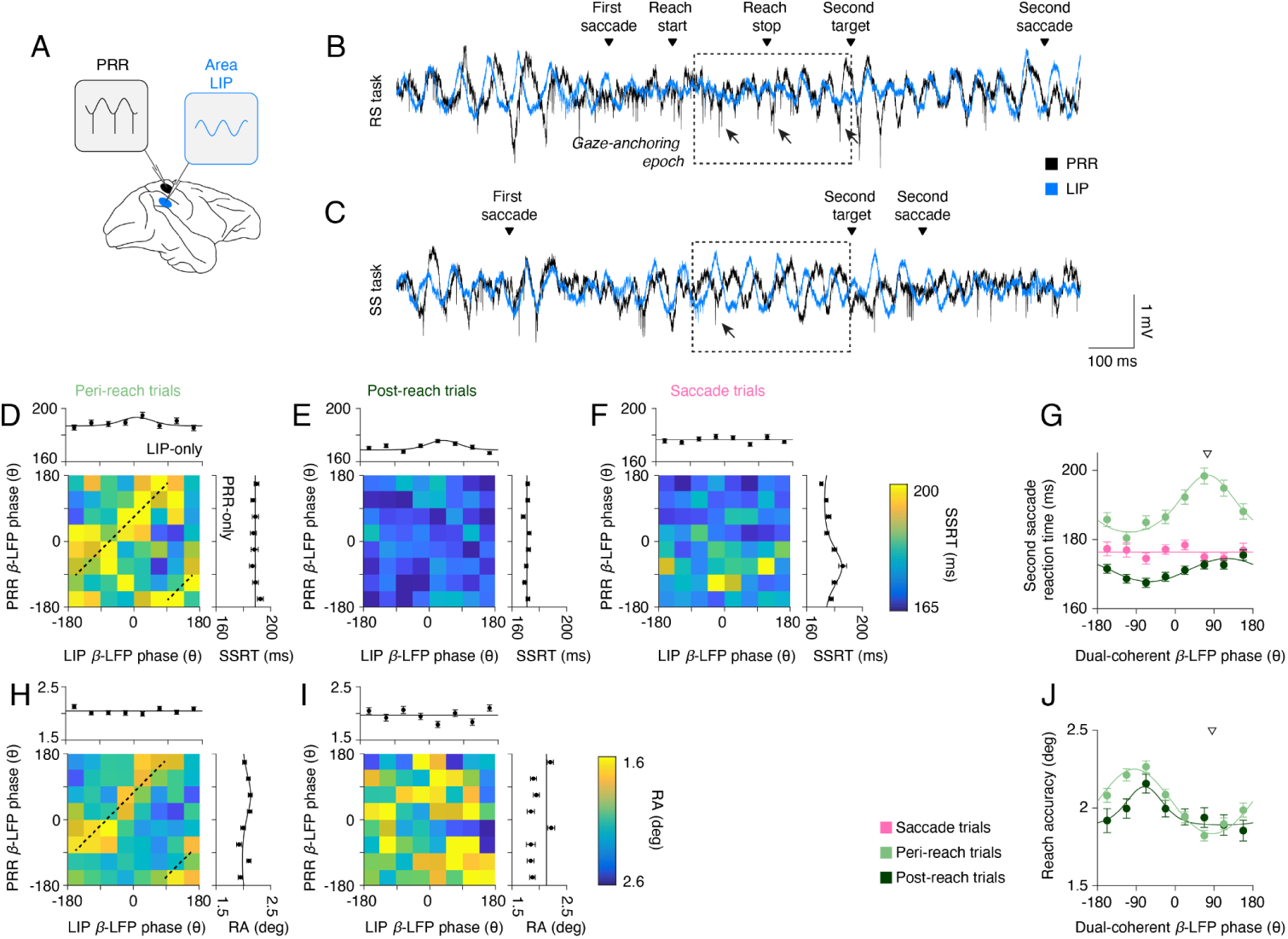
Behavioural performance depends on coherent neural dynamics. (**A**) Schematic showing experimental set up and neural recordings used in subsequent analyses. (**B**) Voltage traces of RS task and (**C**) SS task example trials (PRR: black, LIP: blue). Dashed box indicates gaze-anchoring epoch used as the analysis window for neural data. Arrows indicate example spikes occurring at representative phases in each task. (**D**) Peri-reach, (**E**) post-reach, and (**F**) saccade trials showing mean β-LFP phase in each cortical area (PRR β-LFP phase, y axis; LIP β-LFP phase x axis) and their influence on Second saccade reaction time (SSRT, colorscale). Marginals show SSRT as a function of mean PRR-spike β-LFP phase in each area alone. (**G**) SSRT as a function of dual-coherent β-LFP phase for each trial type (Peri-reach trials, light green; Post-reach trials, dark green; and Saccade trials, pink). Solid lines present changes in SSRT fitted by von Mises function. (**H**) Peri-reach and (**I**) post-reach trials showing mean β-LFP phase in each cortical area (PRR β-LFP phase, y axis; LIP β-LFP phase x axis) and their influence on reach accuracy (RA, colorscale). Marginals show RA as a function of mean PRR-spike β-LFP phase in each area alone. (**J**) RA as a function of dual-coherent β-LFP phase for each trial type, same conventions as in (G). Downward triangles present the mean of the von-Mises fit dual-coherent β-LFP phase on peri-reach trials at maximum SSRT (**G**) or minimum RA (**J**). Dashed lines (D,H) indicate the corresponding dual-coherent phase shown by the downward triangles in (G) and (J), respectively.

We next investigated whether beta-frequency spike-LFP phase predicted trial-by-trial changes in coordinated behaviour. To characterize coherent spiking trial-by-trial, we calculated the beta-frequency phase of LFP activity in area LIP and PRR at times when PRR neurons fired spikes, which we term dual-coherent phase (see **Methods**). We then averaged phase across spiking events during the gaze-anchoring epoc and calculated SSRT and reach accuracy across trials according to the phase of PRR spiking with respect to PRR beta-frequency LFP activity, termed PRR-only phase, and with respect to area LIP beta-frequency LFP activity, termed LIP-only phase.

Analyzing SSRT revealed that variations in PRR spiking with PRR-only phase were not consistent with gaze anchoring. Trial-by-trial changes with PRR-only phase did not predict SSRT on peri-reach or post-reach trials but did predict SSRT on saccade trials (**Fig 2D-F**, PRR-only. peri-reach: p=0.09. post-reach: p=0.40. saccade: p=3.7×10^−12^. von Mises test). Variations of PRR spiking with LIP-only phase were also inconsistent with gaze-anchoring because SSRT was most strongly predicted on post-reach trials (**Fig 2D-F**, LIP-only. peri-reach: p=0.025. post-reach: p=6.4×10^−6^. saccade: p=0.12. von Mises test).

These results demonstrate that changes in PRR spiking with respect to phase in either PRR or area LIP cannot support reach-to-saccade communication during gaze-anchoring. This is because they do not predict variations in performance during peri-reach trials when reach-to-saccade communication is expected due to activity in the reach system. Recent work links multiregional communication to spike timing with respect to the phase of beta-frequency coherence in both regions, termed dual-coherence^35^. Dual-coherent multiregional communication implies that neural interactions between the parietal reach and parietal saccade regions reflect reach spike timing with respect to population activity in both regions. If so, multiregional communication between the reach and saccade systems may occur when beta-frequency coherence has a consistent phase difference across the reach-and-saccade system and may be suppressed at other times.

We computed the dual-coherent phase for each trial by subtracting beta-frequency phase in area LIP from beta-frequency phase in PRR at the time of each PRR spike. We then averaged the SSRT across trials with a similar dual-coherent phase for each task condition. Variations in performance with PRR spike dual-coherent phase were consistent with gaze-anchoring. On peri-reach trials, SSRT significantly varied trial-by-trial with dual-coherent phase and was slowest on trials with a preferred dual-coherent phase of ∼75° **(Fig 2G**, p=2.2×10^−16^, von Mises test, **Methods**). SSRT did not significantly vary with dual-coherent phase on saccade trials when reach-to-saccade was not expected (**Fig 2G**. p=0.5, von Mises test). SSRT on peri-reach trials with non-preferred dual-coherent phases did not increase compared with SSRT on saccade trials (For example, SSRT peri-reach vs saccade trials phase bin centered at -112°, p=0.25, permutation test). SSRT significantly varied with dual-coherent phase on post-reach trials (**Fig 2G**. p=1.40×10^−5^, preferred phase = 120°, von Mises test). However, post-reach SSRT slowing was less pronounced than peri-reach SSRT slowing. On peri-reach trials, SSRTs varied with dual-coherent phase by 17 ms. On post-reach trials, SSRT trials varied with dual-coherent phase by 6 ms.

These data demonstrate that the relationship between SSRT slowing and dual-coherent PRR spike timing is consistent with reach-to-saccade communication on reach trials not saccade trials and consistent with more reach-to-saccade communication on peri-reach trials than on post-reach trials.

We analyzed whether PRR spike timing also predicted trial-by-trial variations in reach accuracy. PRR spiking with LIP-only phase did not predict improved reach accuracy (**Fig 2H,I**. peri-reach: p=0.36; post-reach: p=0.2; von Mises test). Interestingly, PRR spiking with PRR-only phase did predict reach accuracy on peri-reach trials and not post-reach trials (**Fig 2H,I**. peri-reach: p=6.4×10^−3^, post-reach: p = 1, von Mises test). However, reach accuracy varied relatively weakly with PRR-only phase on peri-reach trials (variation in reach accuracy (RA): 0.15° 7% fractional change (ΔRA/mean(RA)). In contrast, reach accuracy significantly and more strongly depended on dual-coherent phase, (**Fig 2J**, variation in RA = 0.45°, 22% fractional change, p=0, von Mises test). On post-reach trials, RA also significantly depended on dual-coherent phase albeit more weakly than on peri-reach trials (**Fig 2J**, variation in RA=0.25°, 12% fractional change, p=4.2×10^−3^, von Mises test).

Taken together, variations of reach accuracy and SSRT with dual-coherent phase were consistent with the presence of a common underlying mechanism of reach-to-saccade communication. On peri-reach trials, reaches were most accurate and SSRT slowest at a similar dual-coherent phase (Reach accuracy phase: 91°. SSRT phase: 75°). Variations of reach accuracy and SSRT with single-region phase were not consistent with a common underlying mechanism of reach-to-saccade communication.

Parametrically-fitting SSRT and RA to phase variations trial-by-trial reinforced these results and showed dual-coherent phase had greater likelihood and less generalization error compared to single-region phase. (**Methods, Extended Data Fig 5**). A non-parametric analysis of SSRT and RA with respect to phase trial-by-trial also emphasized the importance of dual-coherent phase compared with single-region phase (SSRT: dual-coherent phase: resultant = 6.2×10^−3^, p < 10^−6^, LIP-only phase: p = 0.28, PRR-only phase: resultant = 2.3×10^−6^, p = 0.02, RA: dual-coherent phase: resultant = 1.6×10^−4^, p = 0, LIP-only phase: p = 0.33, PRR-only phase: resultant = 5.3×10^−5^, p = 1.6×10^−3^, see **Methods**). The non-parametric analysis was consistent with the absence of reach-to-saccade communication on dual-coherent saccade trials (SSRT: p = 0.89) and consistent with more reach-to-saccade communication on peri-reach trials than on post-reach trials (post-reach SSRT: resultant = 5.7×10^−3^, p = 0. post-reach RA: resultant = 2.1×10^−4^, p = 1.2×10^−3^).

One potential concern is that dual-coherence on peri-reach trials is driven by the evoked LFP phase change due to the onset of the reach. To address this concern, we calculated the PRR-spike dual coherence aligned to reach onset, after explicitly subtracting the evoked LFP response. Reach-onset-aligned PRR spike dual-coherence continued to strongly predict SSRT slowing on peri-reach trials (**Extended Data 6**). Confounds due to the onset of the Go cue were not a concern because the LFP phase calculation during gaze-anchoring epoch rarely overlapped cue delivery (3.7% of peri-reach trials, 0% of post-reach trials and 2.5% of saccade trials).

Additional analyses emphasized the importance of PRR-spike timing with respect to beta-frequency dual-coherent phase. Variations in the phase of beta-frequency LFP phase alone did not predict gaze-anchoring-related SSRT slowing (**Extended Data Fig 7**). Beta-frequency coherence has a period of 50 ms, which implies spike timing changes every quarter-cycle, e.g. 12.5 ms. We therefore jittered PRR spike times on each trial and recalculated dual-coherent phase (see **Methods**). Consistent with 10 ms spike-timing, PRR-spike dual-coherent phase on peri-reach trials predicted SSRT when spike times were jittered by 5 ms or less and no longer predicted SSRT when spike times were jittered by 10 ms or more (**Extended Data Fig 8**).

Gaze anchoring effects were specific to PRR-spike dual coherence in the beta frequency (20 Hz) compared to the gamma frequency (40 Hz, see **Extended Data Fig 9**). Gamma-frequency PRR-spike dual-coherent phase had a small but significant effect on SSRT on peri-reach trials, and size of this effect was significantly smaller than for beta-frequency PRR-spike dual-coherent phase (resultant = 2.9×10^−3^, p = 1×10^−4^ compared to beta-frequency dual-coherent phase, permutation test).

Area LIP spike rate predicted SSRT slowing trial-by-trial (**Fig 1J**). Therefore, we asked whether area LIP spike timing with respect to dual-coherent phase also predicted SSRT slowing. LIP-spike beta-frequency dual-coherent phase had a small but significant correlation with SSRT on peri-reach trials (**Extended Data Fig 10**). However, SSRT varied with PRR-spike dual-coherent phase significantly more than with LIP-spike dual-coherent phase (resultant=1.2×10^−3^, p<10^−6^, permutation test).

These results indicate that specifically PRR spike timing may drive behavioural inhibition during gaze-anchoring to slow SSRT and improve reach accuracy with respect to beta-frequency dual-coherent phase compared with single-region beta-frequency phase, LFP coherence phase, gamma-frequency coherence, and LIP spike timing.

### A reach-to-saccade communication channel

Since spiking in PRR does not generally guide saccades, the manner by which PRR spiking can suppress the second saccade could depend on modulation of a reach-to-saccade communication channel. According to this channel hypothesis, on trials when the reach-to-saccade channel opens, SSRT lengthens because PRR firing is more effective at suppressing saccades. Conversely, on trials when the reach-to-saccade channel closes, SSRT shortens because PRR firing is less effective at suppressing saccades. We therefore analyzed the relationship between PRR spiking and SSRT for evidence of such modulation-state-dependent reach-to-saccade communication.

PRR firing covaried with gaze-anchoring-related increases in SSRT specifically on trials with preferred dual-coherent phase (**Fig 3A**, R=0.15, p=9×10^−4^, Pearson correlation). SSRT did not change when the PRR dual-coherent phase was 180° out-of-phase (non-preferred, R= 0.01, p = 0.82), even when PRR-sender activity increased ten-fold to 30 sp/s. Covariation of combined PRR spiking and modulation state with SSRT was specific to gaze-anchoring and not present during post-reach and saccade trials (**Fig 3B**, post-reach: R=0.01, p=0.85. **Fig 3C**, saccade: R=-0.02, p=0.81, Pearson correlation at preferred phase).

**Figure 3.**
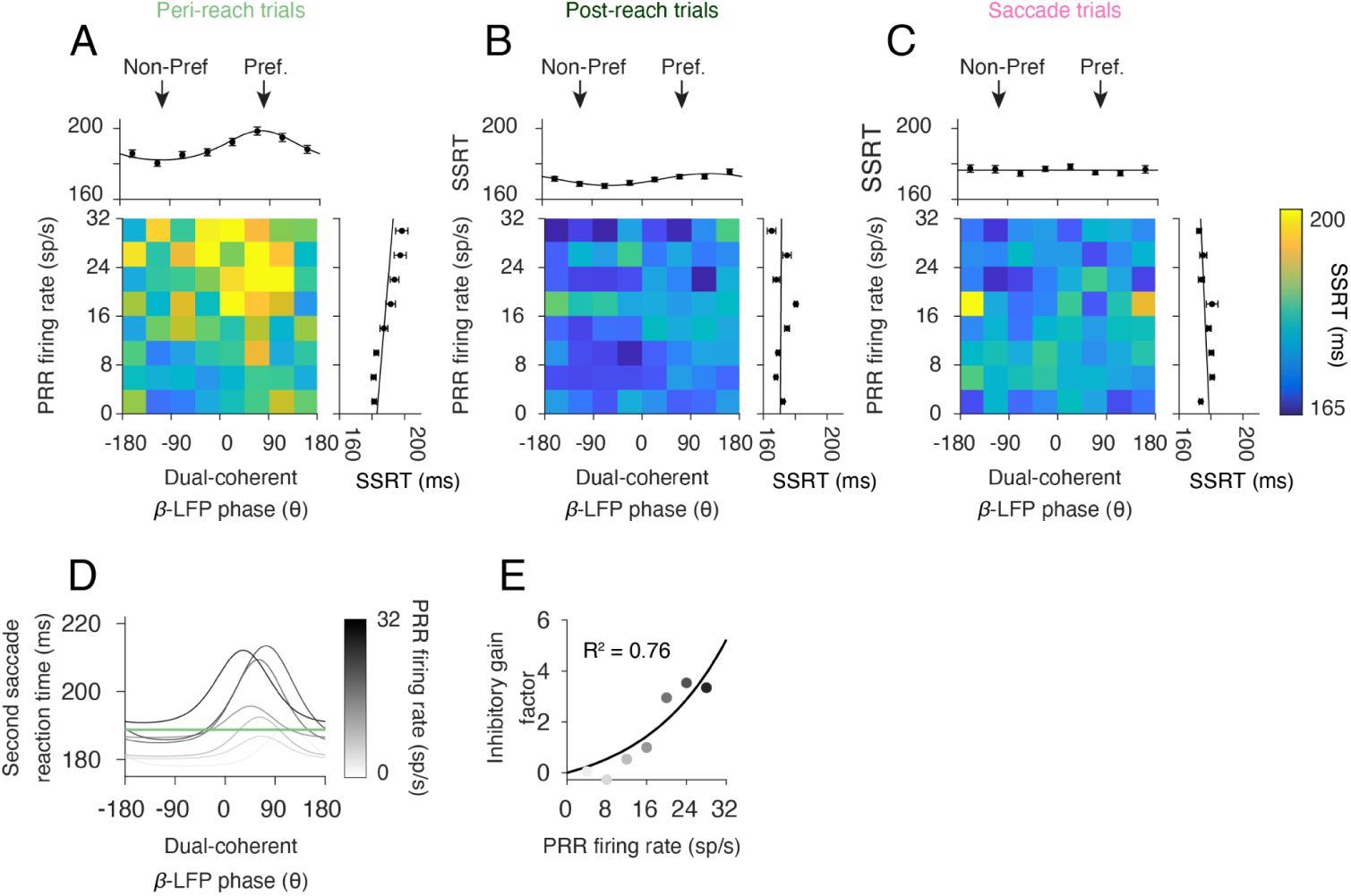
State-dependent inhibitory look-reach communication. (**A**) Peri-reach, (**B**) post-reach, and (**C**) saccade trials showing PRR firing rate (y-axis) and dual-coherent β-LFP phase (x-axis) and their influence on second saccade reaction time (SSRT, colorscale). Marginals show SSRT as a function of PRR fring rate or dual-coherent β-LFP phase alone. Arrows indicate phase bins used for ‘Preferred’ and ‘Non-preferred’ channel states. (**D**) Modulation state functions fit to PRR firing rates bin for peri-reach trials. Horizontal green line indicates mean SSRT for peri-reach trials. (**E**) Inhibitory gain factor function fit to the scaled peaks of the modulation state functions presented in (D) for each PRR firing rate bin.

These data suggest that gaze-anchoring-related increases in SSRT depend on a state-dependent gain. The input drive from PRR is gain-modulated by channel state, e.g. dual-coherent phase. When certain modulation states involve large changes in PRR activity that are effectively compressed and have small gain, the channel is effectively closed. During other modulation states, the same changes in PRR activity can lead to changes with larger gain and the channel is effectively open.

To better understand the relationship between channel gain and channel modulation state in PRR activity, we divided trials into groups according to the level of PRR firing and fit the variation of SSRT with dual-coherent phase on those trials (**Fig 3D**). This analysis revealed a gain mechanism. As PRR firing rate increased, SSRT slowed more on trials where the channel was more open. Plotting the gain factor for different levels of PRR firing revealed a non-linear slowing of SSRT with increases in PRR activity (**Fig 3E**, adjusted R^2^ = 0.76). The state-dependent non-linear gain underlies how the channel can be more open or closed.

Analyzing the relationship between PRR firing and dual-coherent phase revealed that PRR neurons fired the same number of spikes across trials independent of dual-coherent phase during gaze-anchoring (**Extended Data Fig 11**). Since the overall rate of PRR spiking does not depend on dual-coherent phase, the level of PRR firing is not modulated by channel state, which is consistent with PRR fulfilling the role of a sender in this circuit.

### Inhibitory channel modulation predicts LIP suppression

The inhibitory-channel-modulation model explains how gaze-anchoring is controlled by reach-to-saccade communication according to the relationship between PRR firing, dual-coherent phase and SSRT. The model also makes testable predictions about how the saccade system in general, and area LIP firing in particular, should depend on PRR firing and dual-coherent phase. **Figure 4A** illustrates the model and predictions. According to the model, PRR-sender activity varies trial-by-trial and acts as input to the communication channel. The communication channel transforms the PRR input to suppress activity in area LIP from the pre-move epoch to the move epoch, as well as saccade behaviour. Critically, the model predicts that inhibitory modulation operates according to two dissociable components: an **inhibitory gain** function models the inhibitory gain for the PRR firing on that trial, and a **modulation state** function models the state of the dual-coherent phase on that trial (see **Methods**).

**Figure 4.**
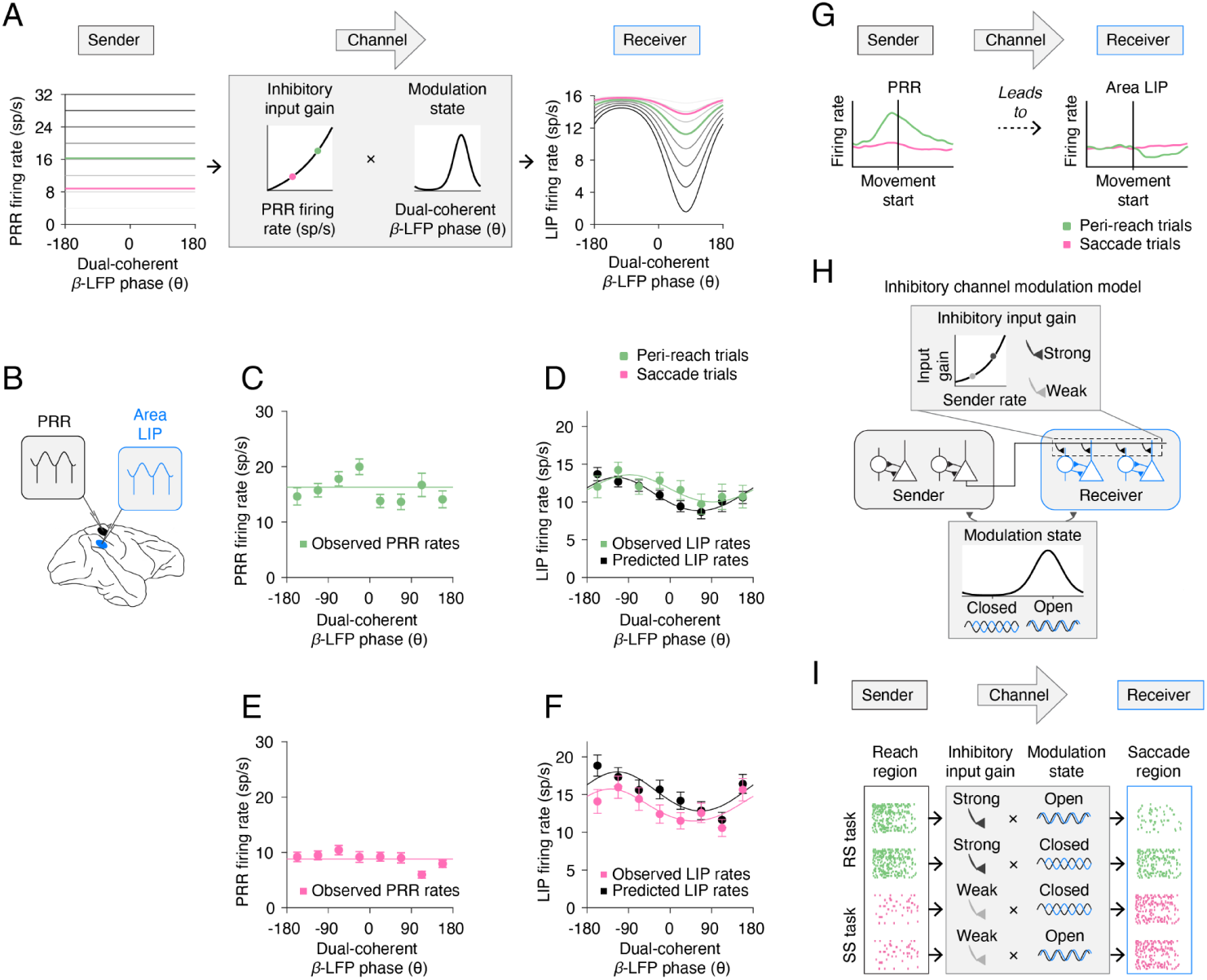
Inhibitory channel modulation model predicts LIP firing rates. (**A**) Inhibitory channel modulation model takes inputs as the PRR firing rate and dual-coherent β-LFP phase. The PRR rate is used to determine the inhibitory input gain and the dual-coherent β -LFP phase is used to determine the modulate state. The product of these two functions makes a prediction about the LIP firing rate on that trial. (**B**) Schematic showing the dual-area neural recordings used to test the model predictions. (**C**) Observed PRR firing rates on peri-reach trials (light green) as a function of dual-coherent β-LFP phase. (**D**) Observed LIP firing rates on peri-reach trials (light green) and predicted LIP firing rates from the model (black) as a function of dual-coherent β-LFP phase. (**E**) Observed PRR firing rates on saccade trials (pink) and (**F**) observed (pink) and predicted (black) LIP firing rates on saccade trials as a function of Dual-coherent β-LFP phase. (**G**) Schematic showing Sender activity in PRR leads to suppressed Receiver activity in area LIP. (**H**) This inhibition is predicted by the inhibitory modulation model which consists of an inhibitory input gain function which ranges from weak to strong and a modulation state function which ranges between open and closed. (**I**) High sender activity during the RS task may lead to strong inhibitory input, which on ‘open’ state trials leads to suppression in the receiver area. Whereas low sender activity during the SS task means inhibitory input gain is weak and does not drive suppression regardless of modulation state.

Because the properties of the saccade system are reflected in saccade behaviour, we model and fit the inhibitory input gain function and the modulation state function using saccade behaviour without directly observing neural activity in the saccade system. The inhibitory input gain and the modulation state are observed and fit using PRR firing and dual-coherent phase from **Fig 3**. Moreover, since area LIP, the receiver, reflects the output of the communication channel that guides saccade behavior, the model predicts area LIP activity. Area LIP firing should be suppressed from the pre-move epoch to the move epoch during the RS task but not the SS task. The amount of suppression should follow the strength of PRR firing, which determines the inhibitory gain, and the dual-coherent phase, which determines the modulation state. In this way, the inhibitory-channel-modulation model predicts suppression of area LIP activity from pre-move epoch before gaze anchoring. The model does not observe area LIP activity during the move epoch that reveals gaze-anchoring itself (see **Methods** for details).

To test the model, we analyzed simultaneously-recorded firing of PRR neurons, LIP neurons, SSRT and PRR dual-coherent phase (32 spatially-selective PRR-LIP neuron pairs; 88 spike-spike-LFP-LFP sessions also featuring simultaneous PRR-LIP LFP recordings; **Methods, Fig 4B**). We specifically measured PRR firing rates and dual-coherent phase to predict simultaneously-recorded LIP firing on each trial. As expected, although PRR firing did not vary with dual-coherent phase (**Fig 4C**), area LIP firing significantly covaried with dual-coherent phase (**Fig 4D**, green, Peri-reach: p=0; von Mises test) and was maximally suppressed at the same preferred phase angle as the SSRT variations with dual-coherent phase (minimum firing rate at 92°, von Mises fit). These features of area LIP firing were predicted by using the observed PRR firing and dual-coherent phase on each trial as input to the model (**Fig 4D**, black).

Finally, we evaluated the relative contributions of the inhibitory input gain, the modulation state and their combination to the predicted area LIP firing rates by evaluating three reduced models (**Methods**). The inhibitory-channel-modulation model best predicted the observed area LIP firing rates during the RS task. Removing the non-linear inhibitory input gain reduced the quality of the prediction more than removing the modulation state function. A linear regression model fit by observing simultaneous recordings of firing rates of PRR and area LIP neurons performed substantially worse, despite being fit by observing area LIP activity during gaze anchoring (mean squared error: inhibitory-channel-modulation model = 175; inhibitory-input gain only = 177; modulation-state only = 185; linear regression = 247; **Methods**). At lower PPR fring rates, as observed in the SS task, the model predicted weak modulation of LIP firing rates (**Fig 4A**). Consistent with this, area LIP firing rates in the SS task were also predicted by using the observed PRR firing and dual-coherent phase on each trial as input to the model (**Fig 4E**,**F**, mean squared error: inhibitory-channel-modulation model = 137; inhibitory-input gain only = 138; modulation-state only = 285; linear regression = 232). Therefore, the results of the modelling suggest PRR firing can specifically influence area LIP firing during gaze-anchoring to coordinate looking and reaching.

## Discussion

Here, we investigate the mechanisms of inhibitory communication during eye-hand coordination. We analyze the activity of individual neurons to establish the behavioural significance of trial-by-trial variations in inhibitory multiregional communication. We show that gaze-anchoring, a naturally-expressed behavioural strategy, may be mediated by a dynamic, state-dependent communication channel. Individual neuron activity in the parietal saccade system (area LIP) is consistent with modulated inhibitory input sent from neurons in the parietal reach system (PRR) to decrease parietal saccade neuron firing and improve behavioural performance (illustrated in **Fig 4G**).

We use behavioral task design to decompose a naturally-expressed behaviour into components necessary to test the mechanisms of multiregional communication by single neurons^36,37^. Gaze-anchoring isolates communication from the reach system to the saccade system. Area LIP activity related to the second saccade could not be responsible for signals in PRR that improve reach accuracy because we presented the second saccade target after the reach and we placed the saccade targets in locations such that area LIP neural response fields for the first and second saccades did not overlap in space. Since gaze-anchoring is a naturally-expressed behaviour, confounding influences due to behavioural training that could alter the underlying mechanisms are absent.

Our results suggest that beta-frequency neural coherence modulates how much reach-related firing suppresses saccade-related firing (**Fig 4H**). During the RS task, reach activity is high, inhibitory input gain is strong and the saccade region is more or less suppressed depending on the modulation state. During the SS task, reach activity is low, inhibitory input gain is weak and the saccade region does not depend on modulation state (**Fig 4I**).

We show that inhibitory communication involves beta coherence and not gamma coherence, complementing work on excitatory communication^29^. Slower saccades and accurate reaches occur with relative phase of ∼75°. Since beta-frequency, 20 Hz, activity has a period of 50 ms, 75° corresponds to a ∼10 ms time difference. This is consistent with the latency for presynaptic PRR spike propagation across U-fibers to area LIP, as well as post-synaptic LIP hyperpolarization due to inhibitory GABA synapses. Previous work also links beta coherence has been linked to GABAergic activity both experimentally^38–41^ and through modelling^38^. Taken together, our results suggest that beta-frequency coherence may specifically engage feedforward inhibition by suppressing synaptic influences from PRR on area LIP across an inhibitory feedforward pathway^42,43^. Note that our results do not imply that LFP activity exerts causal influences on brain function. Interacting populations of neurons may instead exert causal influences that are measured by relative LFP phase.

Our work offers important constraints on theoretical explanations of how multiregional neural population dynamics exert causal effects on behavior. We show that the mechanism of dynamic modulation depends on spike timing at 5-10 ms time-scales. Neural population dynamics at slower 50-100 ms time-scales may explain the gain component but not the modulation component. Therefore, relatively fast firing dynamics are needed in order to explain the mechanisms of behaviorally-relevant communication.

Inhibitory control mechanisms have been implicated in flexible, coordinated behaviour^25,44,45^, visual attention^34,46^ and dual task performance^47,48^. We show that an increase in the reach-related firing of individual neurons in PRR is associated with net suppression of firing by individual neurons in area LIP and the slowed initiation of saccades to visual targets presented unexpectedly at different spatial locations. Net suppression and slowed saccade initiation is consistent with suppressed attentional selection throughout area LIP. Consequently, beta-frequency modulation may allow the reach system to transiently suppress attentional selection in the saccade system. Beta-frequency inhibitory multiregional communication may reflect a general mechanism of inhibitory cognitive control necessary to support flexible behavior.

## Data availability

The datasets generated during the current study are available from the corresponding author on reasonable request.

## Code availability

Matlab code for current study is available on a github repository.

## Acknowledgements

We would like to thank Gerardo Moreno for surgical assistance, Roch Comeau, Stephen Frey and Brian Hynes for custom modifications to the BrainSight system, and Nic Price, Elizabeth Zavitz, Adam Charles and members of the Pesaran lab for helpful feedback. This work was supported, in part, by NIH T32 EY007136 (MAH), ARC DE180100344 (MAH), NSF CAREER Award BCS-0955701 (BP), NEI R01-EY024067 (BP), the Army Research Office (BP), the Simons Foundation (BP), a McKnight Scholar Award (BP), and a Sloan Research Fellowship (BP). MAH and BP conceptualized the project and designed the experiments. MAH performed the experiments. MAH and BP analyzed the data and wrote the manuscript. The authors declare no competing interests.

## Methods

### Experimental Preparation

Two male rhesus monkeys (*Macaca mulatta*) participated in the experiments (Monkey 1, 9.5 kg and Monkey 2, 6.5 kg). Each animal was first implanted with an MRI-compatible head cap under general anesthesia. A structural MRI was obtained with 0.5 mm isotropic voxels and used to guide the placement of a recording chamber over the posterior parietal cortex of the hemisphere contralateral to the reaching arm (Monkey 1: right reaching arm and left hemisphere; Monkey 2: left reaching arm and right hemisphere) in a second surgical procedure. Chamber placement and electrode recording sites were registered to the structural MRI to within 1 mm (BrainSight, Rogue Research). The structural MRIs were also used to estimate recording locations for area LIP and PRR (see **Extended Data Fig 3**). All surgical and animal care procedures were done in accordance with National Institute of Health guidelines and were approved by the New York University Animal Care and Use Committee.

### Behavioural experiments

#### Experimental hardware and software

Eye position was monitored with a video-based eye tracker (I-Scan). Visual stimuli were generated using an array of tristate light-emitting diodes (LEDs, Kingbright, USA) situated directly behind a touch screen (ELO Touchsystems). The LEDs formed a grid with points spaced at 10° intervals. The use of LEDs to present visual stimuli allowed for precise temporal control of stimulus onset and offset. LEDs also ensured that there was no source of background illumination that could influence reach accuracy. Reach accuracy was measured by calculating the Euclidean distance between the target LED and the position of the hand on the touch screen. Trials for which the hand position at reach completion was more than 5° from the target were excluded from further analysis. The visual stimuli were controlled via custom LabVIEW (National Instruments) software executed on a real-time embedded system (NI PXI-8184, National Instruments).

#### Experimental design

Each monkey first performed a center-out saccade task to map the spatial saccade response fields of neurons. On a subset of sessions, each monkey also performed a center-out reach-and-saccade task to map spatial reach response fields of neurons. Each monkey then performed the reach-and-saccade double-step task, RS trials, or the saccade-saccade double-step task, SS trials, to study gaze-anchoring in a manner that was consistent with natural behaviour. On a subset of catch trials (percent of trials: Monkey 1: 15%, [13-18%], Monkey 2: 17%, [15-23%], median, interquartile range), subjects performed only the first step of the double-step tasks with no second target to suppress anticipation. All RS, SS and catch trial conditions were randomly interleaved.

#### Center-out tasks

At the start of each trial, ocular fixation and manual touch were instructed by a green target and a red target, placed centrally side-by-side. The green target indicated the start position for the hand touch, and the red target indicated the start position for the eye. The subject fixated while touching the screen for a variable baseline period of 500-800 ms. In the center-out saccade task a red saccade target would appear in the periphery. In the center-out reach-and-saccade task a yellow saccade target would appear in the periphery. There were eight possible target locations in each task. Each monkey then maintained fixation and touch for a variable delay period of 1000-1500 ms. After the delay period, the central fixation target would extinguish, cueing each monkey to saccade to the target location while maintaining hand position at the initial touch position for the center-out saccade task, or reach-and-saccade to the target location for the center-out reach-and-saccade task.

#### Reach-and-saccade double-step task (RS task)

Initial fixation and touch were again instructed by a red target and a green target, respectively. The initial position was placed 10 degrees to the left (Monkey 1) or right (Monkey 2) of the central target on the horizontal axis, ipsilateral to the recording chamber. Each monkey touched and fixated for a variable baseline of period of 500-800 ms, after a yellow target would appear at the central location. After a variable delay of 1000-1500 ms, the initial touch and fixation were extinguished cueing a reach and saccade to the yellow target. The second saccade target was presented after the reach was completed after an interval of 10-800 ms. The second saccade target was a red LED cueing a saccade alone, presented after the reach was completed after an interval of 10-800 ms and placed either in the response field of the area LIP neuron under study or at an alternative target location also positioned in the contralateral visual field but outside the response field.

#### Saccade-saccade double-step task (SS task)

Initial fixation was cued by a red target 10° away horizontally from the central target and the initial touch was cued by a green target at the central target location. The first saccade target also appeared at the central location, cuing the first saccade toward the hand. As a result, the hand-eye position before the second saccade was identical to that during the RS task. After the baseline period, a red target would appear at the central location. After a variable delay of 500-800 ms, the initial fixation target was extinguished cueing a saccade alone to the central target. As in the RS task, the second target was a red LED cueing a saccade alone. The second target was presented 10-1000 ms after the first saccade.

Overall, visual and oculomotor spatial and temporal contingencies were matched between the two tasks so that the tasks were naturalistic, did not require dedicated training and differed according to whether or not the subject made a reach. In pilot experiments, we also observed that presenting the second saccade target after the first saccade and during the reach resulted in changes in coordinated visual behaviour that altered the timing of the coordinated reach and saccade and led to inconsistent task performance. This was likely due to confusion about the cues, and their interference with ongoing visual processes needed to guide the first movement, such as attention. Since presenting second saccade targets during the reach would require training to ensure consistent task performance, we only studied presentations of the second saccade target after the reach was completed, which both monkeys could perform successfully without the need for additional training.

#### Behavioural database

We collected a database of trials from each monkey for each task (Monkey 1: 10,324 RS task, 8,372 SS task; Monkey 2: 12,840 RS task, 8,452 SS task) across 10 task conditions that were randomly interleaved. This allowed us to analyze the relationship between the latency of the second saccade and the variables of the two tasks in sufficient detail to identify and test multiregional communication. Trials in which saccade and reach reaction times, for both steps, were not within a 100-500 ms window were discarded. This ensures that on all trials analyzed, the subject was neither anticipating nor being inattentive to the targets.

#### SSRT vs second target delay

We compared second saccade reaction time to second target delay from first saccade (both tasks), second target delay from reach completion, reach reaction time, reach duration, and reach reaction time minus saccade reaction time for the first step (RS task only). For presentation, we graph the independent variable in 10 ms bins (**Fig 1C**). Each bin needed a minimum of 20 trials to be included in the analysis, although the average number of trials was usually much greater (Monkey 1: RS task, 198±130 mean±SD; [118:195] interdecile range; SS task, 162±29 mean±SD; [75:387] interdecile range; Monkey 2: RS task, 263±178 mean±SD; [29:220] interdecile range; SS task, 158±63 mean±SD, [49:493] interdecile range).

#### SSRT vs reach accuracy

We measured the association between the second saccade reaction time and the accuracy of the reach by performing linear regression and reporting the slope, statistical significance and correlation coefficient separately for peri-reach trials and post-reach trials (**Fig 1D**, Monkey 1: 3,825 Peri-reach, 2,921 Post-reach trials; Monkey 2: 6,635 Peri-reach, 4,329 Post-reach trials).

### Neurophysiological experiments

*Experimental design:* We performed neuronal recordings during a subset of task conditions used to study behaviour. In the RS task, the second target was either presented 10-300 ms after the reach completion, which we will refer to as **peri-reach trials**, or 500-800 ms after reach completion, which we will refer to as **post-reach trials**. In the SS task, the second target was presented 200-1000 ms after the first saccade to temporally match the second target presentation to that in the RS task accounting for the duration of the reach, which we will refer to as **saccade trials**. On average, the reach was initiated 165±39 ms (Monkey 1, mean±SD; [124:218] interdecile range) or 123±65 ms (Monkey 2, mean±SD; [94:153] interdecile range) after the Go cue with a reach duration of 171±42 ms (Monkey 1, mean±SD, [128:211] interdecile range) or 122±33 ms (Monkey 2, mean±SD, [91:159] interdecile range). We defined the 350 ms time period prior to second target onset as the **gaze-anchoring epoch**.

Neural recordings were made from area LIP and PRR on the lateral and medial banks of the intraparietal sulcus using multiple-electrode microdrives (Double MT, Alpha Omega; **Extended Data Fig 3**). Neurons were recorded within 5-8 mm of the cortical surface. Spiking and LFP activity were recorded with glass-coated tungsten electrodes (Alpha Omega) with impedance 0.7–1.4 MΩ measured at 1 kHz (Bak Electronics). Neural signals were amplified (×10,000; TDT Electronics), digitized at 20 kHz with 12 bits/sample (National Instruments), and continuously streamed to disk during the experiment (custom C and Matlab code). Broadband neural activity was preprocessed to obtain single-unit spike times and LFP activity. All significant differences in firing rates for this study were determined using a random permutation test with 10,000 permutations.

During the experiment, we analyzed the activity of each area LIP neuron recorded in the center-out saccade task to assess spatial selectivity. If the LIP neuron appeared to show spatial selectivity, the double-step tasks were run, including all of the test conditions described above. We placed the second target for the double-step tasks either within the response field for an area LIP neuron being recorded, or at an alternative location in the same visual hemi-field outside the response field. During the experiment, PRR neurons were isolated and recorded regardless of their response properties.

#### Area LIP neuronal database

Area LIP neurons were isolated and mapped for spatial selectivity using a visually-guided, center-out, delayed saccade to eight possible target locations, as described above. After the experiment, if the cell showed a significant increase in activity during the delay period of the center-out task relative to baseline period for a given target (p<0.05, permutation test), the cell was determined to be spatially-selective and that target was labeled as being in the cell’s preferred direction. Each LIP cell was recorded for a minimum of 10 trials in the preferred direction for each task condition (peri-reach, post-reach and saccade trials). If the LIP cell met these two criteria (spatial selectivity in a center-out task and minimum number of trials), the cell was included in the database. Importantly, there were no inclusion or exclusion criteria for the LIP neurons based on neural responses in either of the double-step tasks.

#### PRR neuronal database

After the experiment, we analyzed the activity of each PRR neuron for responses to planning and executing the reach. We only included in the database PRR neurons that contained a significant response during the delay and reach execution periods of the RS task when compared to the baseline epoch of that task. For a minority of PRR neurons, (13/34 neurons) we also confirmed the location of the first movement was in the response field by mapping the response field in the center-out reach-and-saccade task using 4-8 targets. PRR neurons that did not respond to the first movement of the RS task compared to the baseline and PRR neurons with responses to other target locations were excluded from further analysis. Consequently, the first movement of the RS task was in the reach response field of the PRR neurons under study.

#### Firing rate RS/SS task selectivity

We estimated peri-stimulus time histograms with a 20 ms smoothing window. We defined a task selectivity index that measured the fractional difference (MFD) in firing rate between the RS and SS task trials ((RS-SS)/SS) during the gaze-anchoring epoch. We tested for significant differences between the RS and SS task trials by comparing the measured task selectivity index with a null distribution of task selectivity indices when randomly permuting the RS and SS task labels on each trial. Results presented in **Fig 1E,F,G,H**.

#### Firing rate vs SSRT

We measured the association between the second saccade reaction time and the firing rate of area LIP and PRR neurons by performing linear regression and reporting the slope, statistical significance and correlation coefficient separately for peri-reach trials, post-reach trials and saccade trials. Results presented in **Fig 1I,J**.

#### LFP phase

We subtracted the mean LFP response from each trial to suppress the influence of responses evoked by the stimuli and responses. We then band-pass filtered the LFP at 20 Hz to study beta-frequency activity and at 40 Hz to study gamma-frequency activity. Band-pass filtering was performed with multitaper methods (T= 200 ms, W= 5 Hz; ^49^). Due to variability in the timing of the coordinated reach and saccade and the temporal smoothing necessary to resolve band-limited LFP phase, the peri-reach interval and not the post-reach interval potentially includes reach execution, reach preparation and the coordinated saccade.

#### Dual-coherent LFP phase

For each trial, we measured the phase of LFP activity at the time of the spiking activity by calculating the spike times within the analysis window and computing the phase of band-pass filtered LFP activity at these times. The mean phase was calculated for each trial by calculating the circular mean across all spikes within the analysis window. We refer to this value as the spike-triggered dual-coherent LFP phase. For spike-LFP-LFP sessions, we calculated the spike-triggered LFP phase for each spike-LFP pair (ϕ_*PRR*_ and ϕ_*LIP*_) and then the circular distance between the two phases, which we refer to as the dual-coherent phase (ϕ_*Dual*_):

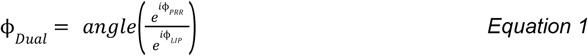

The circular statistics toolbox in Matlab (The Mathworks) was used to perform statistical tests (Bergens, 2009). Results presented in **Fig 2, Extended Data Figs 4, 6-10**.

Phase analysis also allowed us to analyze trial-by-trial variations between the neural responses and behavioural effects such as the reaction time for the second saccade and accuracy of the reach, described below.

#### LFP phase difference and SSRT

For each trial, we measured the effect of LFP phase alone on SSRT. For each area, we calculated the circular mean of the phase of the LFP across the analysis window. We then calculated the circular distance between the two phases in each area using *Equation 1* above. Results presented in **Extended Data Fig 7**.

#### Dual-coherent LFP phase vs SSRT - parametric approach

We modeled the SSRT from peri-reach, post-reach and saccade trials using a von Mises fit in which SSRT varies across trials according to a gamma distribution Γ(*k*, θ)with constant scale, *k*, and a rate, θ, that depends on the phase, ϕ, on that trial during the last 350 ms preceding the onset of the second target according to a von Mises function, θ = *A* + *B exp*(κ *cos*(ϕ− μ)). We defined three different versions of the model each containing the same number of parameters where phase was set by PRR-only phase, LIP-only phase or dual-coherent phase. The gamma scale parameter, *k*, and the von Mises fit parameters, *A, B*, κ, μ, were estimated using maximum likelihood. The null hypothesis was that SSRT varied across trials according to a gamma distribution with constant scale and rate parameters and did not vary with phase. For each version of the model, the likelihood was maximized using the function mle in Matlab (Mathworks). We fit parameters using a two-step procedure. In step 1, we initialized parameters based on heuristics derived from the SSRT vs phase tuning curve. The offset, *A*, was initialized at the minimum of the tuning curve.

The magnitude, *B*, was initialized using the range of the tuning curve. The preferred phase, μ, was initialized at the phase with the maximum of the tuning curve. The dispersion, κ, was initialized at 0.5 based on visual inspection of the tuning curves. The scale, *k*, was also initialized at 20 based on visual inspection of the SSRT distributions. When tuning was weaker, the fits based on initializing based on heuristics became trapped in local minima. In such cases, we pursued step 2. In step 2, we generated surrogate data sets by jittering the SSRT observations by adding a random value less than 1% of the original data and refitting the data using the same heuristics as before. We then used the parameter fits obtained from the surrogate data to initialize the optimization for the original data and repeated the optimization based on these initial conditions. We tested for significance of the von Mises fit for each version of the model against the null hypothesis using a likelihood-ratio test. We selected between the models based on dual-coherent phase, LIP-only phase and PRR-only phase based on the difference in the maximized log likelihood according to Akaike Information Criterion (AIC). For each model, we also estimated and compared the generalization error using k-fold cross validation with 10 folds. Results presented in **Fig 2D-G, Extended Data Fig 5A**,**B**.

To test the dependence of dual-coherent phase on spike timing, we repeated the analysis described above after jittering the spiketimes on each trial according to a Gaussian distribution with standard deviation 2 ms, 5 ms,10 ms or 20 ms. Results presented in **Extended Data Fig 8**.

#### Dual-coherent LFP phase vs reach accuracy - parametric approach

We modeled the accuracy of the reach on peri-reach and post-reach trials according to a von Mises fit in which accuracy varies across trials according to a gamma distribution Γ(*k*, θ) with constant scale, *k*, and a rate, θ, that depends on the phase, ϕ, on that trial during the last 350 ms preceding the onset of the second target according to a von Mises function,θ = *A* + *B exp*(κ *cos*(ϕ− μ)). We defined three different versions of the model each containing the same number of parameters where phase was set by PRR-only phase, LIP-only phase or dual-coherent phase. For each model, the gamma scale parameter, *k*, and the von Mises fit parameters, *A, B*, κ, μ, were estimated using maximum likelihood. The null hypothesis was that reach accuracy varied across trials according to a gamma distribution with constant scale and rate parameters and did not vary with phase. The likelihood was maximized using the function mle in Matlab (Mathworks) using the same 2-step procedure as detailed above for SSRT. We tested for significance of the von Mises fit for each version of the model against the null hypothesis using a likelihood-ratio test. We selected between the models based on dual-coherent phase, LIP-only phase and PRR-only phase based on the difference in the maximized log likelihood according to Akaike Information Criterion (AIC). For each model, we also estimated the generalization error using k-fold cross validation with 10 folds. Results presented in **Fig 2H-J, Extended Data Fig 5C**,**D**.

#### Reach accuracy and SSRT vs phase - non-parametric approach

We performed a non-parametric test of the effects of phase on behavioral performance, reach accuracy and SSRT. For each metric, we computed the resultant vector using eight equally-spaced and sized phase bins. To determine significance, we performed a permutation test by permuting the phase on each trial and recalculating the resultant vector (10,000 permutations).

#### Two-sample non-parametric phase tuning

We used a non-parametric test of the effects of phase on behavioral performance to compare resultant vectors across different phase measurements (for example, beta-LFP vs gamma-LFP dual coherence). We performed the permutation test by calculating the difference in resultant vectors and comparing to the resultant when permuting the phases across populations (10,000 permutations). Results presented in **Extended Data Fig 9**,**10**.

#### Reach-start aligned analysis

We analyzed the relationship between LFP phase, SSRT and reach accuracy during a 350 ms time epoch aligned to the start of the reach. The reach-start analysis window extends from 200 ms before the start of the reach until 150 ms after the start of the reach. Since the reach duration was typically 100-200 ms (see **Extended Fig Data 2**), the reach-start analysis window spans the reach execution period. This interval was chosen to be close in time to the gaze-anchoring window while avoiding confounding influences due to presentation of the Go cue and the second saccade target. Earlier time intervals included the onset of the Go cue while later time intervals included the onset of the second saccade target. Results presented in **Extended Data Fig 6**.

#### Dual-coherent LFP phase, firing rate and SSRT

We measured the association between the SSRT and the firing rate of PRR neurons for trials grouped by spike-triggered phase by performing linear regression. We report the statistical significance of the preferred and null phase bins. Results presented in **Fig 3 A-C**.

#### Dual-coherent LFP phase vs firing rate

We modeled the firing rate on peri-reach, post-reach and saccade trials according to a von Mises fit in which spike count varies across trials according to a Poisson distribution with rate, λ, that depends on the phase, ϕ, on that trial during the last 350 ms preceding the onset of the second target according to a von Mises function, λ = *A* + *B exp*(κ *cos*(ϕ− μ)). The von Mises fit parameters, *A, B*, κ, μ, were estimated using maximum likelihood. The null hypothesis was that spike count across trials varied according to a Poisson distribution with constant rate parameter. The likelihood for each model was maximized using the function mle in Matlab (Mathworks). We tested for significance of the von Mises fit against the null hypothesis using a likelihood-ratio test. Results presented in **Extended Data Fig 11, Fig 4C-F**.

#### Inhibitory channel modulation model

We modeled the firing rate of LIP neurons (*Rate*_*LIP*_) trial-by-trial as a function of PRR spike rate (*Rate* _*PRR*_) and dual-coherent β-LFP phase (ϕ_*Dual*_) according to an inhibitory channel modulation model. The model operates according to two functions (see **Fig 4A**): **inhibitory input gain** function models the inhibitory gain (*I*_*t*_) based on the PRR firing on that trial, and a **modulation state** function models the modulation state (*M*_*t*_) based on the dual-coherent β-LFP phase on that trial:

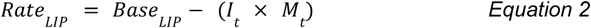

The inhibitory input gain function and modulation state function were fit by binning trials according to PRR spike rate and fitting a von Mises function to the SSRTs in each bin according to dual-coherent phase (see **Fig 3A**). Trials were grouped in increments of 8 spikes/sec. The inhibition scale factor for each firing rate bin was measured as the weighted difference between the peak of the von Mises function fit to that bin and the mean SSRT across all peri reach trials (see **Fig 3D**). The inhibitory gain on each trial (*I*_*t*_) was defined according to an exponential function consisting of two parameters (α, β) and the PRR firing rate on that trial (*Rate*_*PRR*_):

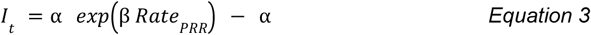

The parameters of the inhibitory input gain function (α, β) were fit using the scale factors of the von Mises fit to each firing rate bin (see **Fig 3E**). The modulation state for each trial was defined according to a von Mises distribution with two parameters (κ, μ) and the dual-coherent β-LFP phase on that trial (ϕ_*Dual*_):

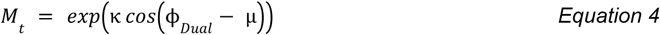

The parameters of the modulation state function (κ, μ) were calculated from the average of the von Mises parameters fit across bins of PRR firing rates and SSRTs (κ = 1. 5, μ = 65 ; see **Fig 3D**). We describe the goodness of fit (R^2^) using the adjusted R^2^ value, which is the ratio of the sum of squared error to the sum of squared total, scaled to account for the number of observations and the number of predictors.

Importantly, LIP spike rates were not used in fitting either the inhibitory gain function or the modulation state function. The inhibitory channel model only depends on LIP spiking activity for a base rate starting point for the model on each trial. The LIP base rate (*Base* _*LIP*_) on each trial (*t*) was defined by the average LIP firing rate recorded before firing rate suppression is observed 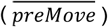 and difference between the LIP firing rate prior to movement onset (*preMove*) and before the onset of the second target (*preTarg*) on each trial such that:

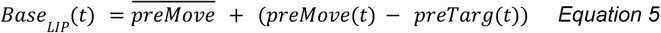

We characterized the performance of the model by calculating the mean squared error (MSE) between the observed LIP firing rate on each trial and the predicted LIP firing rate on each trial according to the model. For comparison, we also calculated the MSE for an input gain function only model, which did not include the modulation state function, a modulation state only model, which did not include the inhibitory input gain function, and a linear regression model. The linear regression, unlike the other models, was fit using the observed LIP firing rates on each trial.

