## Extended Data for "Modulation of inhibitory communication coordinates looking and reaching"

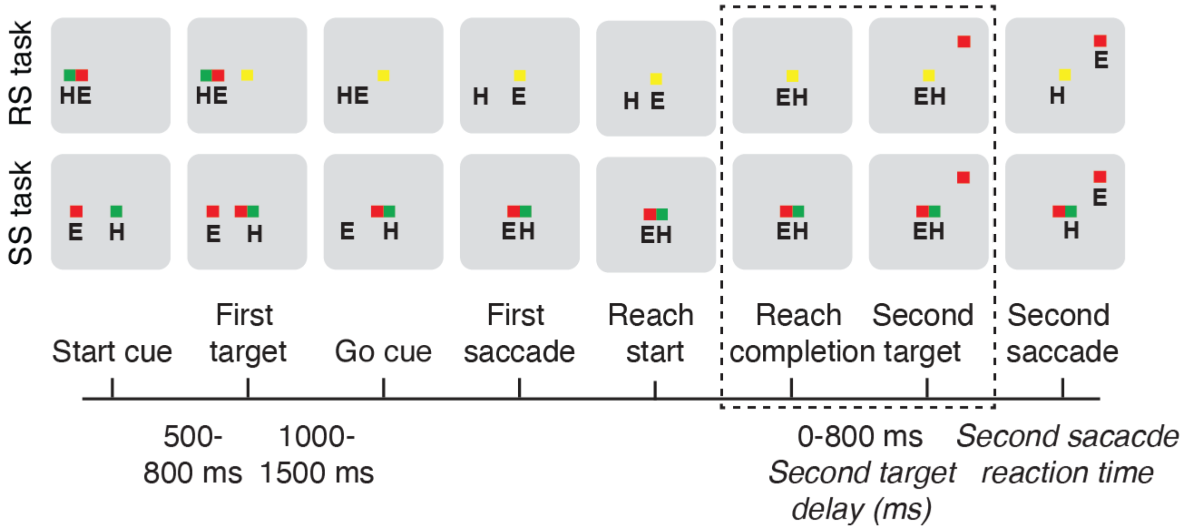


**Extended Data Fig 1.** Coordinated and independent movement tasks. Reach and saccade double-step task (RS) and Saccade double-step task (SS), indicating hand (H) and eye (E) position at each epoch. Dashed lines indicate period of gaze-anchoring in the RS task, and temporally matched epochs in the SS task.

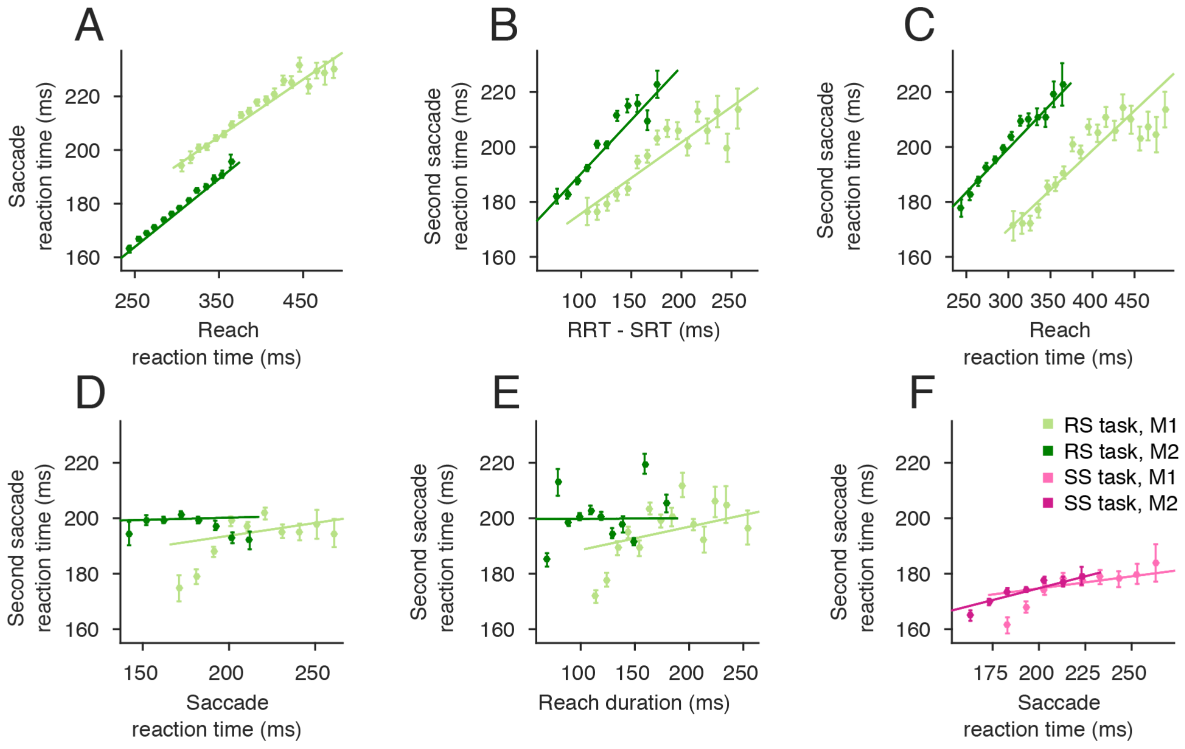


**Extended Data Fig 2.** The slowing of the second saccade reaction time (SSRT) was tied to the coordinated movement. (**A**) The coordination of the first movement was established by the strong correlation between the reaction times for the reach and saccade in the first movement (Monkey 1 (M1): R=0.34, slope=0.21 ms/ms, p=3x$10^{-139}$, Monkey 2 (M2): R=0.45, slope=0.25 ms/ms, p=0, Pearson pairwise linear correlation). (**B**) SSRT correlated with the difference between the reaction times of the reach and saccade in the first movement (M1: R=0.20, slope=0.26 ms/ms, p=2x$10^{-49}$, M2: R=0.22, slope=0.39 ms/ms, p=2x$10^{-78}$, Pearson pairwise linear correlation). (**C**) Specifically, SSRT correlated with the reaction time of the reach (M1: R=0.23, slope=0.29 ms/ms, p=4x$10^{-63}$, M2: R=0.20, slope=0.32 ms/ms, p=2x$10^{-65}$, Pearson pairwise linear correlation). (**D**) The SSRT was not dependent on the reaction time of the saccade in the RS task (M1: R=0.05, slope=0.09 ms/ms, p=8x$10^{-4}$, M2: R=0.006, slope=0.02 ms/ms, p=0.63, Pearson pairwise linear correlation) (**E**) SSRT did not depend on the duration of the reach (M1: R=0.07, slope=0.08 ms/ms, p=1x$10^{-6}$, M2: R=0.001, slope=0.002 ms/ms, p=0.91, Pearson pairwise linear correlation). (**F**) SSRT only weakly correlated with the SRT in the SS task (M1: R=0.06, slope=0.08 ms/ms, p=1x$10^{-3}$, M2: R=0.12, slope=0.17 ms/ms, p=7x$10^{-13}$, Pearson pairwise linear correlation). Therefore, the slowing of the SSRT was tied to coordinated movement, and primarily the timing of the reach.

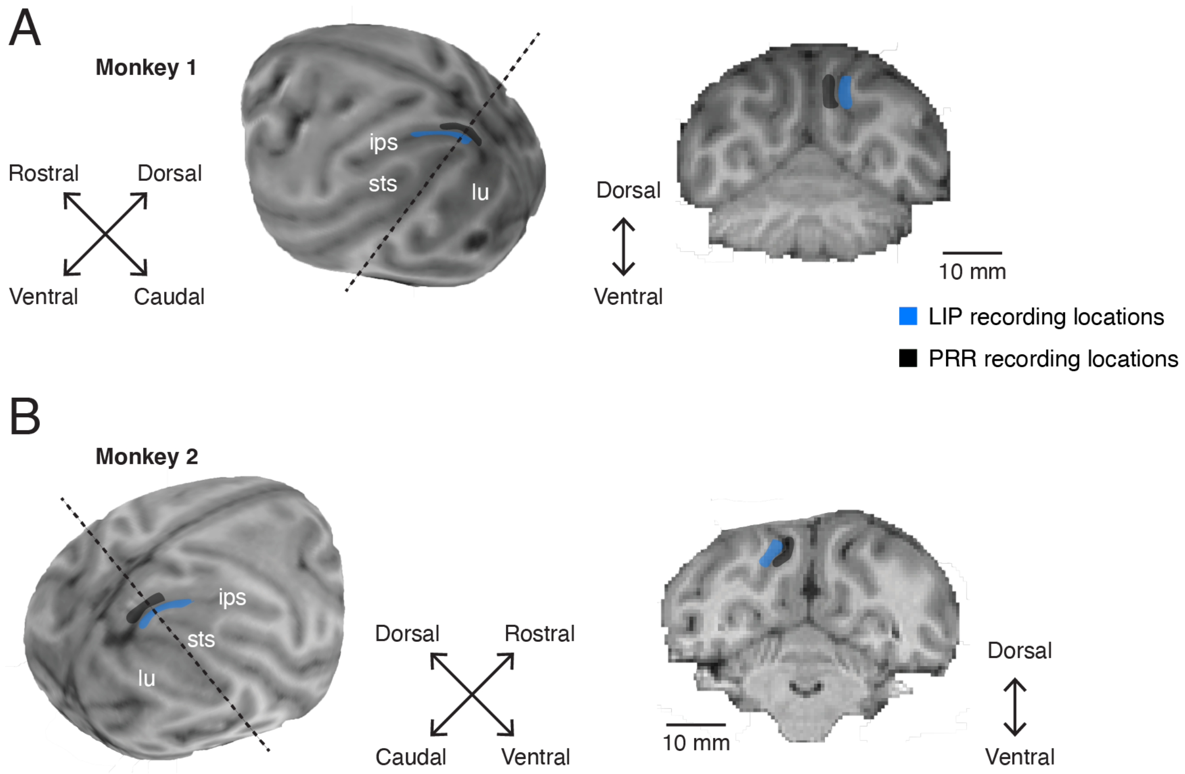


**Extended Data Fig 3.** Anatomical locations of recordings. Recording chambers were placed over the posterior parietal cortex of the hemisphere contralateral to the reaching arm (**A**) Whole brain MRI reconstructions and example coronal slice from Monkey 1 and (**B**) Monkey 2. Chamber placement and electrode recording sites were registered to the structural MRI (BrainSight, Rogue Research). Recording regions for area LIP (blue) and PRR (black) are indicated by the shaded regions. Dashed lines indicate the plane of example coronal sections shown. Key sulcal landmarks, intraparietal sulcus (ips), lunate sulcus (lu) and superior temporal sulcus (sts), are also indicated.

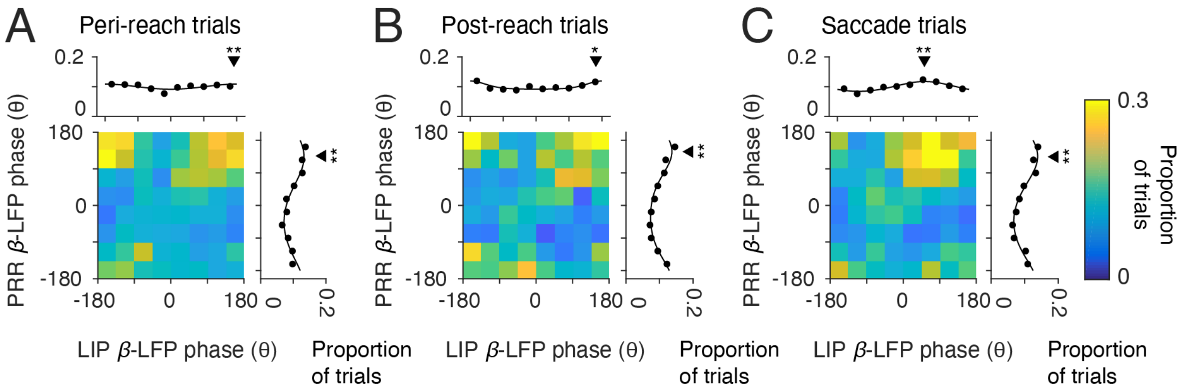


**Extended Data Fig 4.** Phase of spike-β-LFP coherence in each cortical area. (**A**) Peri-reach, (**B**) Post-reach, and (**C**) Saccade trials showing spike-LFP coherence between PRR spiking and PRR LFP phase (y-axis) and LIP LFP phase (x-axis) in the beta-band (β, 20 Hz, colorscale: proportion of trials). Marginals show the proportion of trials as a function of phase in each area. **Peri-reach:** PRR β-LFP $p=2x10^{-49}$, mean phase=136±75°; LIP β -LFP $p=8x10^{-5}$, mean phase=172±79°, **Post-reach:** PRR β-LFP $p=4x10^{-57}$, mean phase=149±74°, LIP β-LFP p=0.02, mean phase=164±79°, **Saccade trials:** PRR β-LFP $p=2x10^{-52}$, mean phase=135±73°, LIP β-LFP $p=2x10^{-52}$, mean phase=59±78°, Rayleigh's test of non-uniformity, circular mean±SD phase). Black triangles indicate mean phase, stars indicate that the distribution is non-uniform (one star, p< 0.05; two stars, p<0.01).

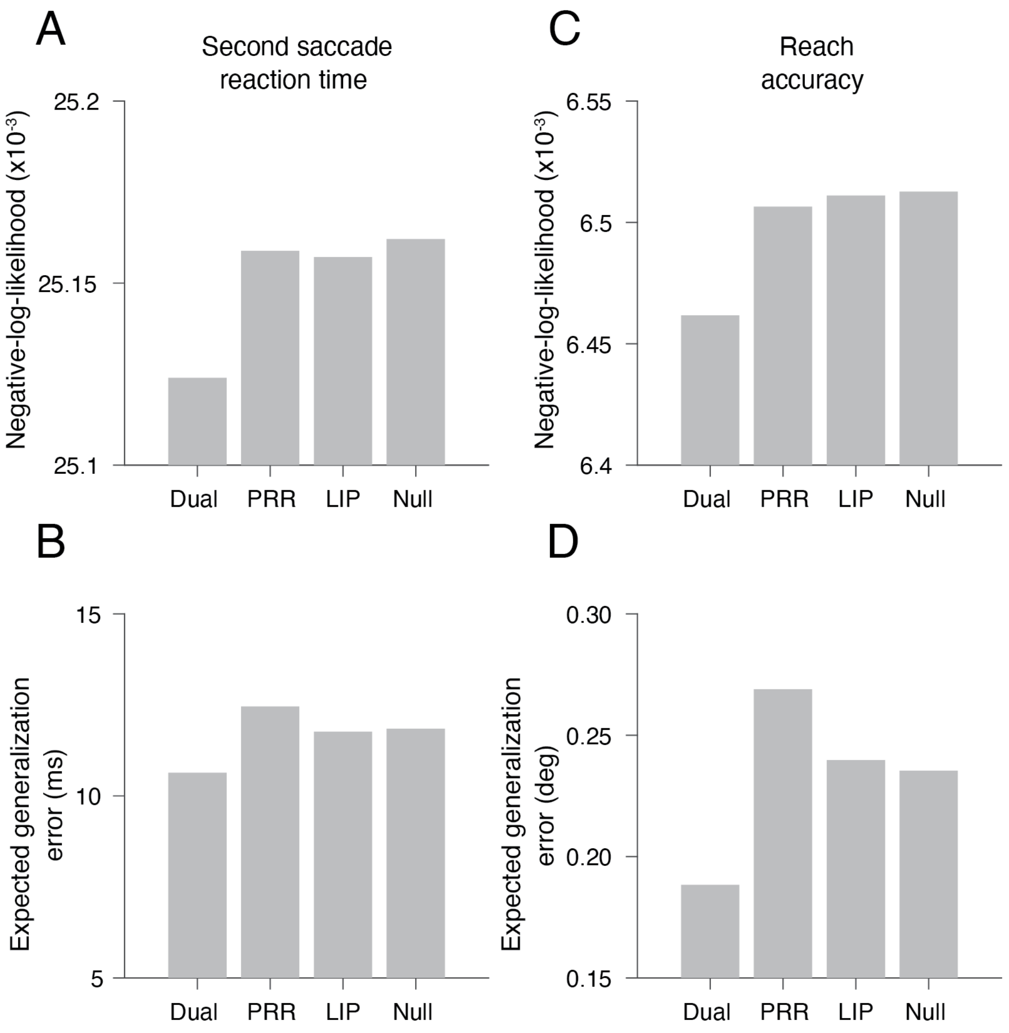


**Extended Data Fig 5.** Negative-log-likelihood and generalization errors for model fits. **Second saccade reaction time: (A)** Dual-phase negative-log-likelihood (NLL)= 25124; PRR-only phase NLL=25158; LIP-only phase NLL = 25157; Null NLL = 25162; $\Delta NLL_{Dual\text{-}PRR}$ = 35; $\Delta NLL_{Dual\text{-}LIP}$ = 33, AIC test; **(B)** Expected generalization error: Dual-coherent: 10.6 ms, R= 0.10; PRR-only: 12.5 ms, R= -0.05; LIP-only: 11.8 ms, R= 0.01; Null: 11.8 ms, where R=1-(SSE_model/SSE_null); **Reach accuracy: (C)** Dual-phase negative-log-likelihood (NLL)= 6462; PRR-only phase NLL=6507; LIP-only phase NLL= 6511; Null NLL = 6512; $\Delta NLL_{Dual\text{-}PRR}$ = 45; $\Delta NLL_{Dual\text{-}LIP}$ = 49, AIC test; **(D)** Expected generalization error: Dual-coherent: 0.18 deg, R= 0.20, PRR-only: 0.27 deg, R = -0.14, LIP-only: 0.24 deg, R = -0.02, Null: 0.24 deg).

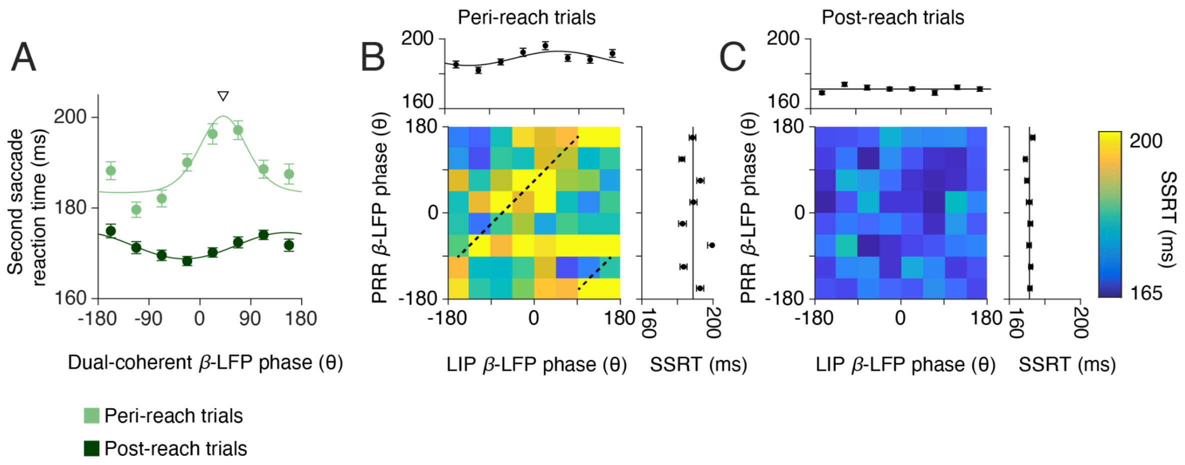


**Extended Data Fig 6.** Dual coherent β-LFP phase aligned to reach onset. (**A**) Second saccade reaction time (SSRT) as a function of dual-coherent β-LFP phase for each RS task trial type (Peri-reach: light green. Post-reach: dark green) during the gaze anchoring epoch when aligned to reach onset, instead of second target onset. Solid lines present changes in SSRT fitted by von Mises function (peri-reach: $p = 0$, preferred phase = 41°. post-reach: $p = 6 \times 10^{-5}$, preferred phase = 153°, von Mises test). Downward triangle presents the mean of the von-Mises fit dual-coherent β-LFP phase at maximum SSRT on peri-reach trials. (**B-C**) Phase of spike-β-LFP coherence in each cortical area (PRR β-LFP coherence, y axis; LIP β-LFP coherence, x axis) and influence on SSRT (colorscale). Marginals show SSRT as a function of β-LFP phase coherence in each area alone (Peri-reach: PRR-only $p = 0.53$, LIP-only $p = 2 \times 10^{-3}$, preferred phase = 48°. Post-reach: PRR-only $p = 0.53$, LIP-only $p = 0.48$, von Mises test) . Dashed lines (**B**) indicate the corresponding dual-coherent phase shown by the downward triangle in (**A**).

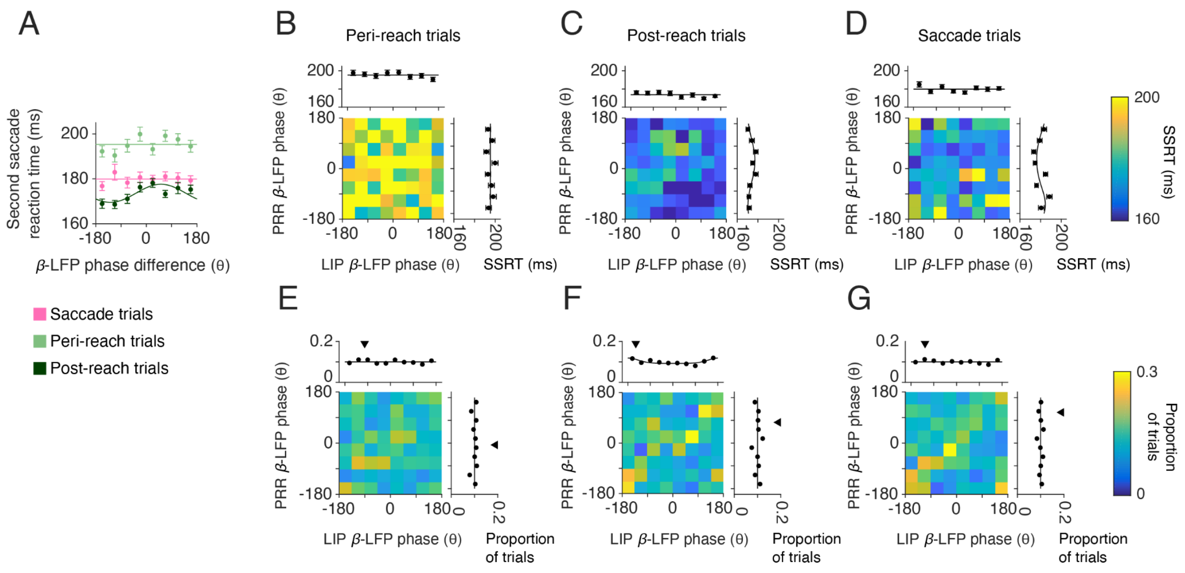


**Extended Data Fig 7.** β-LFP phase difference alone did not predict gaze anchoring. The circular mean β-LFP phase was taken across the gaze-anchoring epoch, irrespective of spike timing. (**A**) Difference in mean phase of the β-LFP (20 Hz) across cortical areas for each task trial type (Saccade: pink. Peri-reach: light green. Post-reach: dark green). Solid lines present changes in SSRT fitted by von Mises function (Peri-reach: $p = 0.23$. Post-reach: $p = 1x10^{-3}$, preferred phase = 53°. Saccade: $p = 0.83$. von Mises test). (**B-D**) Mean β-LFP phase in each cortical area (PRR β-LFP phase, y axis; LIP β-LFP phase x axis) and influence on SSRT (colorscale). Marginals show SSRT as a function of mean β -LFP phase in each area alone (Peri-reach: PRR-only $p = 0.89$, LIP-only $p = 0.24$. Post-reach: PRR-only $p = 8x10^{-3}$, preferred phase = 32° LIP-only $p = 0.20$. Saccade: PRR-only $p = 8x10^{-4}$, preferred phase = -143; LIP-only $p = 0.12$. von Mises test). (**E-G**) β -LFP phase in PRR (y-axis) and LIP (x-axis, colorscale: proportion of trials). Downward triangles show the circular mean phase.

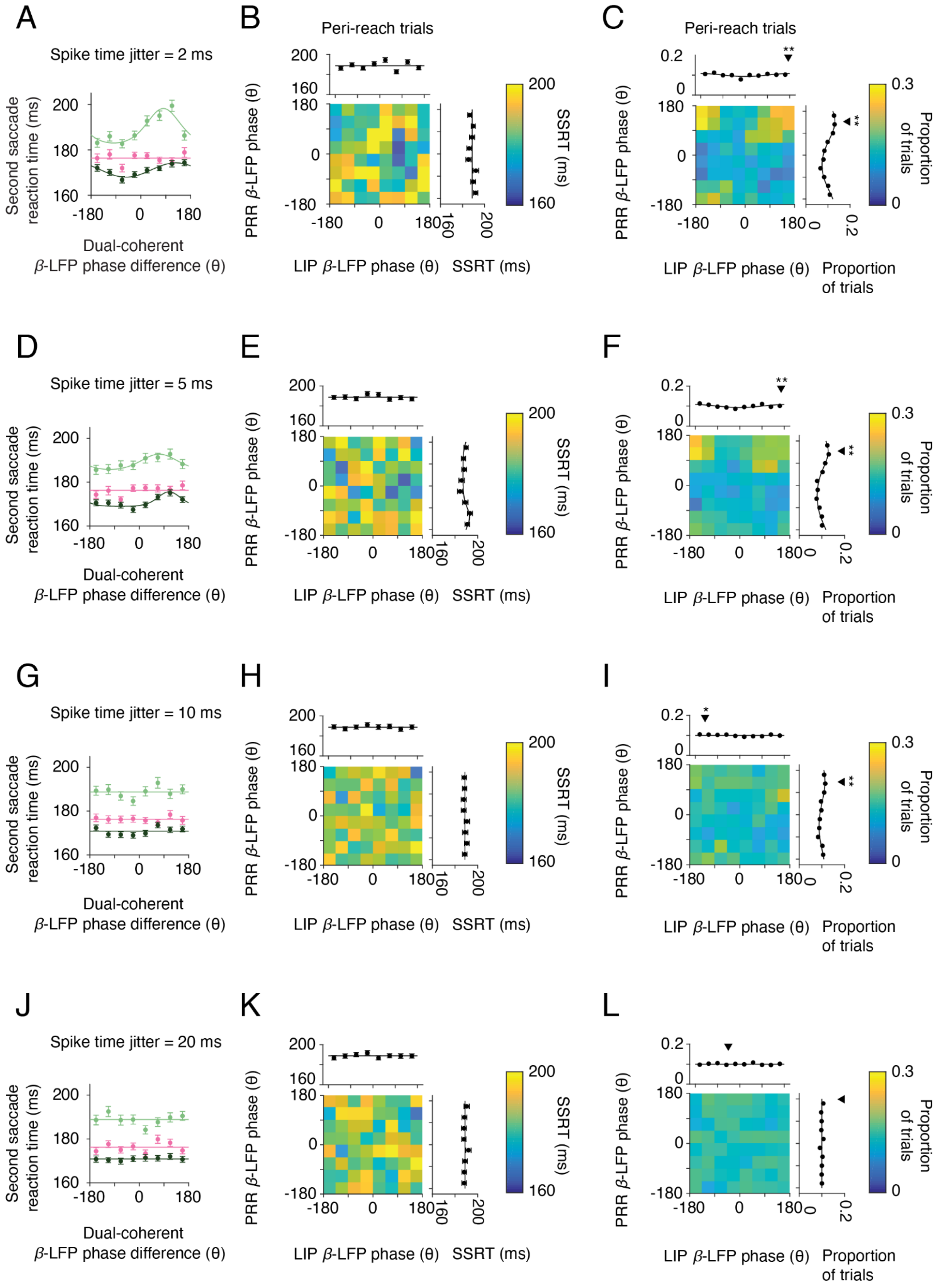


**Extended Data Fig 8.** Spike timing dependence of dual coherent β-LFP phase. For each trial, spike times were jittered according to a Gaussian distribution with standard deviation

(**A-C**) 2 ms, (**D-F**) 5 ms, (**G-I**) 10 ms and (**J-L**) 20 ms. Second saccade reaction time (SSRT) as a function of dual-coherent β-LFP phase for each task trial type (Saccade, pink; Peri-reach, light green; Post-reach, dark green) during the gaze anchoring epoch was recomputed with the jittered spike times. Solid lines present changes in SSRT fitted by von Mises function (**A,** Peri-reach: $p = 1x10^{-13}$, preferred phase = 82°. Post-reach: $p = 5x10^{-5}$, preferred phase = 126°. Saccade: p = 0.68. **D,** Peri-reach: $p = 3x10^{-3}$, preferred phase = 73°. Post-reach: $p = 7x10^{-5}$, preferred phase = 108°. Saccade: p = 0.23. **G,** Peri-reach: p = 0.22. Post-reach: p = 0.29. Saccade: p = 0.90. **J,** Peri-reach: p = 0.11. Post-reach: p = 1. Saccade: p = 0.38. von Mises test). For peri-reach trials, the phase of spike-β-LFP coherence in each cortical area were computed for the jittered spike times (PRR β-LFP coherence, y axis; LIP β-LFP coherence x axis) and influence on SSRT (colorscale). Marginals show SSRT as a function of β-LFP phase coherence in each area alone (**B,** PRR-only p = 0.10, LIP-only, p = 0.10, **E,** PRR-only p = 0.04, preferred phase = -127°, LIP-only, p = 0.47, **H,** PRR-only p = 0.68, LIP-only, p = 0.63, **K**, PRR-only p = 0.52, LIP-only, p = 0.71, von Mises test). Spike-LFP coherence between PRR spiking and each cortical area alone for jittered spike times on peri-reach trials (PRR LFP phase, y-axis, LIP LFP phase x-axis, colorscale: proportion of trials). Marginals show the proportion of trials as a function of phase in each area (**C,** PRR $p = 2x10^{-47}$, mean = 137°, LIP $p = 8x10^{-4}$, mean = 175°, **F,** PRR $p = 2x10^{-31}$, mean = 139°, LIP $p = 1x10^{-3}$, mean = 167°, **I,** PRR $p = 3x10^{-10}$, mean = 132°, LIP p = 0.04, mean = -140°, **L,** PRR p = 0.84, LIP p = 0.79, Rayleigh's test of non-uniformity, circular mean phase). Black triangles indicate mean phase, stars indicate that the distribution is non-uniform (one star, p< 0.05; two stars, p<0.01).

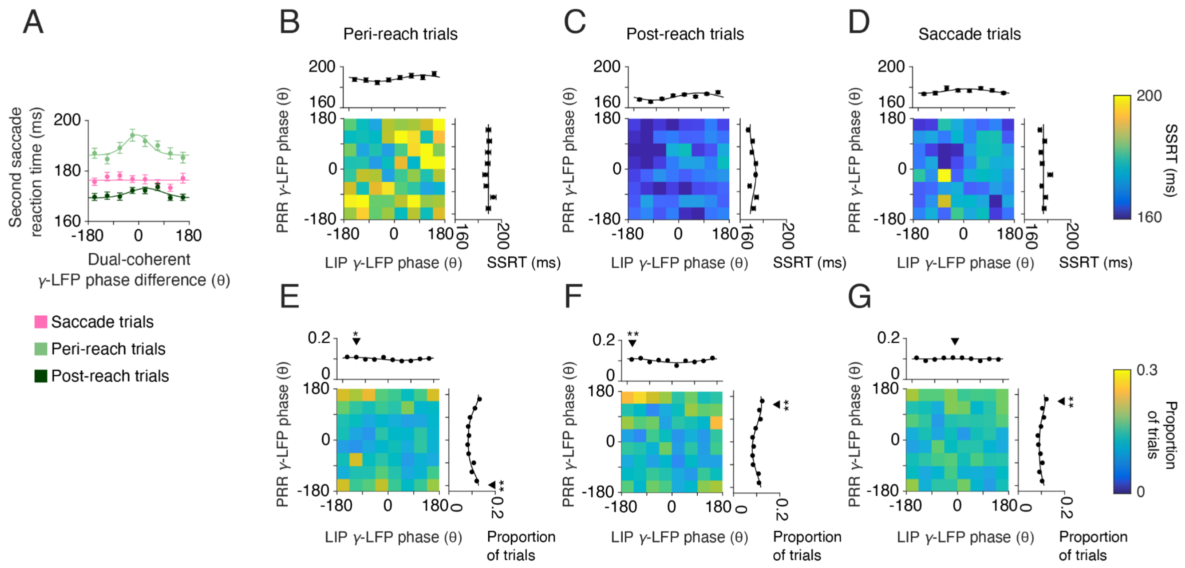


**Extended Data Fig 9.** Dual coherent γ-LFP phase. Dual-coherent phase was calculated between PRR spiking and the γ-LFP phase (40 Hz) in each cortical area. (**A**) Second saccade reaction time (SSRT) as a function of dual-coherent γ-LFP phase for each RS task trial type (Peri-reach = light green; Post-reach = dark green). Solid lines present changes in SSRT fitted by von Mises function (Peri-reach: $p = 8x10^{-4}$, preferred phase = -6°. Post-reach: p = 0.04, preferred phase = 26°. Saccade: p = 0.12 von Mises test). (**B-D**) Mean γ-LFP phase in each cortical area (PRR γ-LFP phase, y axis; LIP γ-LFP phase x axis) and their influence on SSRT (colorscale). Marginals show SSRT as a function of mean γ-LFP phase in each area alone (Peri-reach: PRR-only p = 0.55, LIP-only p = 0.02, preferred phase = 112°. Post-reach: PRR-only p = 0.02, preferred phase = -25°. LIP-only $p = 4x10^{-6}$, preferred phase = 78°. Saccade: PRR-only p = 0.13, LIP-only p = 0.04, preferred phase = 17°, von Mises test). (**E-G**) γ-LFP phase in PRR (y-axis) and LIP (x-axis, colorscale: proportion of trials). Marginals show the proportion of trials as a function of phase in each area (Peri-reach: PRR $p = 4x10^{-23}$, mean = -177°, LIP p = 0.02, mean = -125°. Post-reach: PRR $p = 2x10^{-6}$, mean = 148°, LIP $p = 3x10^{-4}$, mean = -159°. Saccade: PRR $p = 1x10^{-4}$, mean = 153°, LIP p = 0.33, Rayleigh's test of non-uniformity). Black triangles indicate mean phase, stars indicate that the distribution is non-uniform (one star, p< 0.05; two stars, p<0.01).

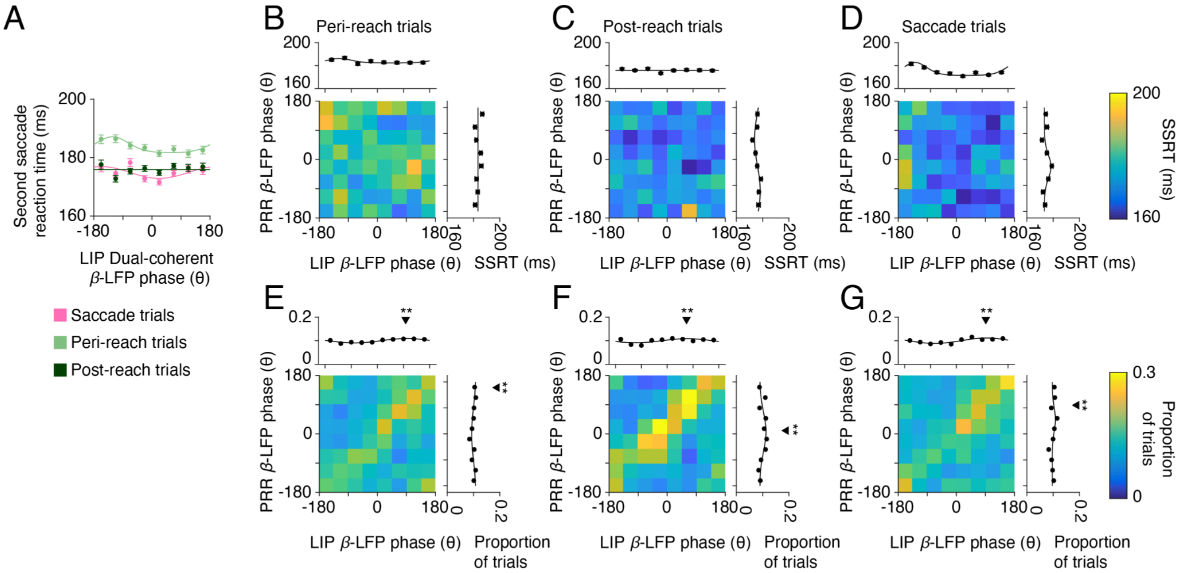


**Extended Data Fig 10.** LIP-spike dual coherent β-LFP phase. Dual-coherent phase was calculated between LIP spiking and the β-LFP phase (20 Hz) in each cortical area. (**A**) Second saccade reaction time (SSRT) as a function of LIP dual-coherent β-LFP phase for each RS task trial type (peri-reach trials, light green; post-reach trials, dark green) and SS task trials (saccade trials). Solid lines present changes in SSRT fitted by von Mises function (peri-reach: $p = 1x10^{-4}$, preferred phase = -122°. post-reach: p = 0.21. saccade: $p = 1x10^{-3}$, preferred phase = -157°. von Mises test). (**B-D**) Mean LIP-spike β-LFP phase in each cortical area (PRR β-LFP phase, y axis; LIP β-LFP phase x axis) and their influence on SSRT (colorscale). Marginals show SSRT as a function of mean LIP-spike β-LFP phase in each area alone (Peri-reach: PRR-only p = 0.23. LIP-only p = 0.49. Post-reach: PRR-only $p = 6x10^{-4}$, preferred phase = 109°. LIP-only p = 0.48. Saccade: PRR-only $p = 5x10^{-5}$ preferred phase = -25°. LIP-only p = 0, preferred phase = -142°, von Mises test). (**E-G**) LIP-spike β-LFP phase in PRR (y-axis) and LIP (x-axis, colorscale: proportion of trials). Marginals show the proportion of trials as a function of phase in each area (Peri-reach: PRR $p = 2x10^{-5}$, mean = 160°, LIP $p = 1x10^{-7}$, mean = 97°. Post-reach: PRR $p = 2x10^{-5}$, mean = 11°, LIP $p = 5x10^{-4}$, mean = 67°. Saccade: PRR $p = 2x10^{-5}$, mean = 100°, LIP $p = 4x10^{-12}$, mean = 103°, Rayleigh's test of non-uniformity, circular mean). Black triangles indicate mean phase, stars indicate that the distribution is non-uniform (one star, $p < 0.05$; two stars, $p < 0.01$).

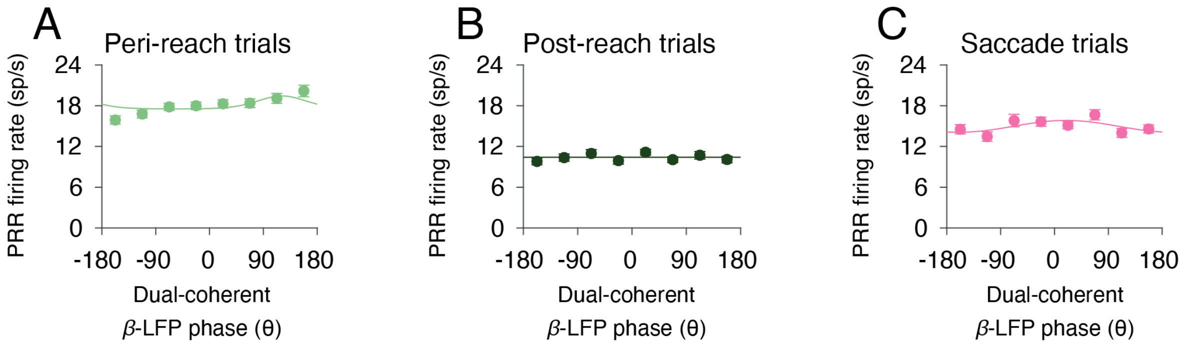


**Extended Data Fig 11.** Dual-coherent β-LFP phase is weakly correlated with PRR firing rate. PRR firing rate and a function of dual-coherent β-LFP phase for each (**A**) Peri-reach, (**B**) Post-reach, and (**C**) Saccade trials. Solid lines present changes in SSRT fitted by von Mises function (Peri-reach: $p = 0$, preferred phase = -121°. Post-reach: $p = 0.23$. Saccade: $p = 0$, preferred phase = 19°, von Mises test).
